# Common genetic variation impacts stress response in the brain

**DOI:** 10.1101/2023.12.27.573459

**Authors:** Carina Seah, Rebecca Signer, Michael Deans, Heather Bader, Tom Rusielewicz, Emily M. Hicks, Hannah Young, Alanna Cote, Kayla Townsley, Changxin Xu, Christopher J. Hunter, Barry McCarthy, Jordan Goldberg, Saunil Dobariya, Paul E. Holtzherimer, Keith A. Young, NYSCF Global Stem Cell Array® Team, Traumatic Stress Brain Research Group, Scott A. Noggle, John H. Krystal, Daniel Paull, Matthew J. Girgenti, Rachel Yehuda, Kristen J. Brennand, Laura M. Huckins

**Affiliations:** Department of Genetics and Genomic Sciences, Icahn School of Medicine at Mount Sinai, New York, NY; Departments of Psychiatry and Genetics, Division of Molecular Psychiatry, Yale University School of Medicine, New Haven, CT; James J. Peters Veterans Affairs Medical Center, Bronx, NY, USA; The New York Stem Cell Foundation Research Institute; Nash Family Department of Neuroscience, Friedman Brain Institute, Icahn School of Medicine at Mount Sinai, New York, NY; Department of Psychiatry, Yale School of Medicine, New Haven CT, USA; US Department of Veterans Affairs National Center for PTSD; Department of Psychiatry, Geisel School of Medicine at Dartmouth, Lebanon, NH 03756, USA; Department of Psychiatry and Behavioral Sciences, Texas A&M College of Medicine, Bryan, Tex.; Central Texas Veterans Health Care System, Temple, Tex.; Department of Psychiatry Center for the Study of Traumatic Stress Uniformed Services, University of the Health Sciences Bethesda MD.

## Abstract

To explain why individuals exposed to identical stressors experience divergent clinical outcomes, we determine how molecular encoding of stress modifies genetic risk for brain disorders. Analysis of post-mortem brain (n=304) revealed 8557 stress-interactive expression quantitative trait loci (eQTLs) that dysregulate expression of 915 eGenes in response to stress, and lie in stress-related transcription factor binding sites. Response to stress is robust across experimental paradigms: up to 50% of stress-interactive eGenes validate in glucocorticoid treated hiPSC-derived neurons (n=39 donors). Stress-interactive eGenes show brain region- and cell type-specificity, and, in post-mortem brain, implicate glial and endothelial mechanisms. Stress dysregulates long-term expression of disorder risk genes in a genotype-dependent manner; stress-interactive transcriptomic imputation uncovered 139 novel genes conferring brain disorder risk only in the context of traumatic stress. Molecular stress-encoding explains individualized responses to traumatic stress; incorporating trauma into genomic studies of brain disorders is likely to improve diagnosis, prognosis, and drug discovery.

**Graphical Abstract:** 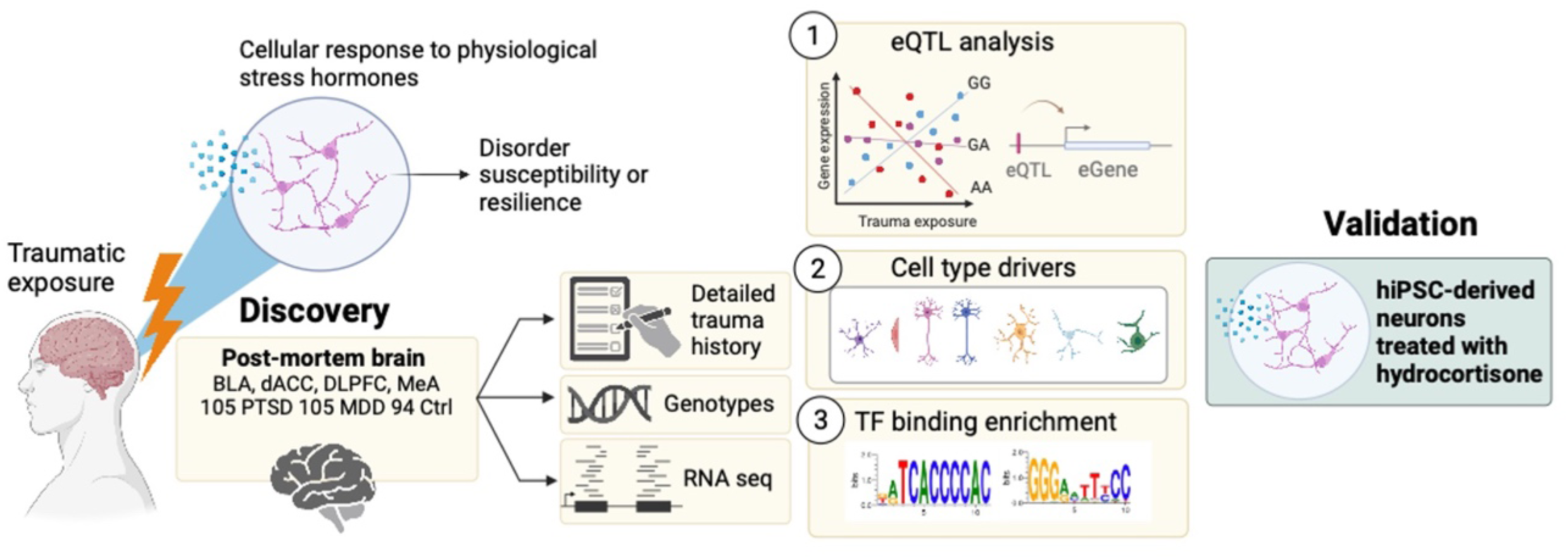

## INTRODUCTION

Traumatic stress is associated with significant physical and psychological comorbidities^1–4^, and increases the risk for and severity of many psychiatric and non-psychiatric medical disorders^5–10^. Given that only some individuals who experience trauma will ultimately develop psychiatric disorders^11^, a long-standing hypothesis is that stress susceptibility may be genetically encoded^12^. The effects of trauma are long-lasting and detectable months or years after exposure^13^; this biological encoding may occur through regulation of gene expression^14–21^. Genetic variation is widely thought to moderate the molecular encoding of stress, likely via a multitude of molecular and cellular alterations including epigenetic modifications of gene expression and downstream effects on glucocorticoid function^22^. For example, polymorphisms in neurotransmitter receptors^23^, metabolizers^24^ and *FKBP5*^25–28^ modify the impact of traumatic stress on psychiatric disorder incidence^29,30^. This can be modelled *in vitro* using human induced pluripotent stem cell (hiPSC)-derived glutamatergic neurons (iGLUTs). For example, iGLUTs from combat-exposed veterans with post-traumatic stress disorder (PTSD) exhibited hyper-responsive glucocorticoid (hCort)-elicited transcriptional signatures relative to those from combat-exposed veterans without PTSD^31^. This diagnosis-dependent transcriptomic response to glucocorticoids raises the question of whether these intrinsic individual differences in stress encoding are mediated by genetic variation. Put simply, how does the variation among individual genomes yield differential stress response?

To answer this requires careful examination of how genomes and environmental factors interact, and in particular how genotype x environment interactions shape higher order biology. Examples of cell-type-specific^32,33^, sex-specific^34–36^, developmental stage-specific^37,38^, and drug exposure-specific^39^ genetic regulation of gene expression abound; however, studies of genotype x stress interactions have progressed more slowly due to difficulties in quantifying stress in genetic and post-mortem brain cohorts. Consequently, it is likely that we have not yet identified many of the genes that confer risk for psychiatric disorders only in the context of extreme stress.

Although human neurons show differential acute response to hCort by diagnosis, leveraging this to predict individualized physiological response to trauma is more complex. Whereas *in vitro* experiments facilitate controlled addition of a single stressor (here, hCort) at a defined dose and duration, modelling trauma in the human brain requires delineation of trauma type, degree, chronicity, and years prior to death, and as a result the impact of accumulated trauma on the human brain is unclear. Likewise, while *in vitro* studies can focus on a single cell type (here, iGLUTs), human brain studies incorporate the effect of dynamic interactions between circuits and cells in the brain. Despite their power for precise comparisons, it is not yet clear how well human neurons *in vitro* model trauma response in the human brain. Thus, for the first time, we investigate if and how human genetic variation confers differential encoding of stress-induced transcriptional phenotypes, examining in parallel post-mortem brains (n=304 donors, with and without known trauma exposure, in tissues from the cortex^40–43^ and amygdala^41–45^) and hiPSC-neurons (n=39 donors, with and without hCort treatment: iGLUTs and iGABAs).

We set out to demonstrate how trauma may dysregulate gene expression in a genotype-specific manner (e.g., trauma-eQTLs) in the post-mortem brain and test whether these are analogous to responses observed *in vitro (*e.g., hCort-eQTLs). To the extent that there is a high degree of overlap between these two measures (e.g., stress-eQTLs), we can reasonably conclude that information obtained from these two disparate analyses yields reliable and functionally meaningful information about genetic contributions to stress responsivity, the development of stress induced psychopathology, and possibly resilience. Modelling these genotype-stress interactions and their impacts on gene expression moreover allows us to examine the potential impacts of stress on a variety of neuropsychiatric disorders. By developing machine-learning models that predict stress-dynamic gene expression on an individual level, we ask whether we can identify new genes associated with these disorders, effectively translating existing ‘static’ case-control genome wide association study results to dynamic insights into genotype-dependent response to stress.

## METHODS

### Human trauma-exposed post-mortem brain cohort

304 post-mortem brain samples (94 control, 105 PTSD, 105 MDD; 136 trauma-exposed; 168 without known trauma exposure) were collected as part of an existing study^42^ (**S. Figure 1A, B**).

A retrospective clinical diagnostic review was conducted on every brain donor, consisting of telephone screening, macroscopic and microscopic neuropathological examinations, autopsy and forensic investigative data, two sources of toxicology data, extensive psychiatric treatment, substance abuse treatment, and medical record reviews, and family informant interviews (i.e., next-of-kin could be recontacted and was agreeable to phone contact, which included the PTSD Checklist (i.e., PCL-5 and/or the MINI). A history of traumatic exposure including exposure to military combat, physical abuse, sexual abuse, emotional abuse, and/or other traumas were obtained as part of the telephone screening, records reviews, and/or PCL-5. A board-certified psychiatrist with expertise in PTSD reviewed every case to rate presence/absence of PTSD symptoms. All data were compiled into a comprehensive psychiatric narrative summary that was reviewed by two board-certified psychiatrists, in order to arrive at lifetime DSM-5 psychiatric diagnoses (including substance use disorders/intoxication) and medical diagnoses. Non-psychiatric healthy controls were free from psychiatric and substance use diagnoses, and their toxicological data was negative for drugs of abuse.

Cumulative trauma burden was quantified by adding up instances of reported traumatic exposure, as previously defined^46–48^. Briefly, traumas included in this analysis included sexual abuse, physical abuse, neglect, witnessing trauma, combat or occupational traumas, assault, and natural disasters. Cumulative trauma was coded independently by three investigators and finally determined by consensus.

### hiPSC-derived neuron cohort

A total of 46 donors (n=23 PTSD, n=23 Control) (**Table S1)** were recruited as previously described. Briefly, participants in this study were combat-exposed veterans with and without PTSD who provided written informed consent (VA HS no. YEH-16-03 and ISMMS HS no. 15-00886) and from whom a viable blood and/or fibroblast sample was obtained and sufficient RNA for genome-wide expression analyses was extracted. All participants experienced deployment to active military combat zones and reported a DSM-IV criterion A combat trauma. Individuals with and without PTSD did not have significant differences in childhood or pre-deployment trauma, deployment number or cumulative duration. Participants underwent psychological evaluation using the Structured Clinical Interview for DSM-5 (SCID) and the Clinician Administered PTSD Scale (CAPS) for determination of PTSD diagnosis and severity. Eligibility criteria and thresholds were based on CAPS for DSM-IV; PTSD(+) had a current CAPS-IV total score above 40 (frequency plus intensity), whereas PTSD(–) participants were combat-exposed veterans with a total score below 40.

### Automated generation of hiPSC-derived ASCL1- and DLX2-induced GABAergic neurons

GABAergic neurons (iGABA) were generated from hiPSCs using high-throughput automated differentiations, in three batches, as previously described^49^ with some modifications. hiPSCs were single-cell passaged after a 20-min dissociation with Accutase (STEMCELL Technologies) at 37 °C and 5% CO2. A total of 1 million cells per well were plated in 12-well Cultrex-coated (R&D Systems, catalog no. 3434-010-02) tissue culture plates (Corning Costar) in PSC Feeder Free Medium (Thermo Fisher Scientific, catalog no. A14577SA) with 1 µM thiazovivin (Sigma-Aldrich, catalog no. SML1045). Lentivirus (generated by ALSTEM) carrying TetO-Ascl1-puro (Addgene, catalog no. 97329), Teto-DLX2-hygro (Addgene catalog no. 97330) and FUdeltaGW-rtTA (Addgene, catalog no. 19780) was diluted to a multiplicity of infection of one each (1 million genome counts of each vector per transduction) in 100 µl DPBS, no calcium, no magnesium (Thermo Fisher Scientific) and added directly after cell seeding. After 24 h, the medium was exchanged (-1.1 mL +1 mL) with Neural Induction Medium (NIM) comprising a 50:50 mix of DMEM/F12 and Neurobasal, with 1× B27 plus vitamin A, 1× N2, 1× Glutamax (Thermo Fisher Scientific) and 1 µM doxycycline hyclate (Sigma-Aldrich). After 24 h, the medium was removed and NIM was added with doxycycline plus 5 µg ml–1 puromycin and 250 µg ml–1 hygromycin B (Thermo Fisher Scientific) (NIM selection medium). Daily medium exchanges were performed with NIM selection medium for 5 days post induction. On day 6 after induction, cells were passaged by incubating with Accutase for 45 min at 37 °C and 5% CO2. A series of 96-well plates (PerkinElmer CellCarrier Ultra) were coated with 0.1% polyethylenimine (Sigma-Aldrich, catalog no. 408727) in 0.1 M borate buffer pH 8.4 for 30 min at room temperature, washed five times with water and prefilled with 100 µl per well of neural coating medium comprising Brainphys medium (STEMCELL Technologies) with 1× B27 plus vitamin A, 1 µM thiazovivin, 5 µg ml–1 puromycin, 250 µM dibutyryl cAMP (dbcAMP, Sigma-Aldrich), 40 ng ml–1 brain-derived neurotrophic factor (BDNF, R&D Systems), 40 ng ml–1 glial cell line-derived neurotrophic factor (GDNF, R&D Systems), 200 µM ascorbic acid (Sigma-Aldrich) and 10 µg ml–1 natural mouse laminin (Sigma-Aldrich). A sample of cells were stained with 10 µg ml–1 Hoechst plus 1:500 acridine orange/propidium iodide solution and counted on an Opera Phenix confocal microscope (PerkinElmer). A total of 100,000 cells per well were seeded into neural coating medium-filled 96-well plates in 100 µl per well of neural medium comprising Brainphys medium with 1× B27 plus vitamin A, 1 µM thiazovivin, 250 µM dbcAMP, 40 ng ml–1 BDNF, 40 ng ml–1 GDNF, 200 µM ascorbic acid and 1 µg ml–1 natural mouse laminin. At 24 h after seeding, medium was exchanged for neural selection medium comprising Brainphys medium with 1× B27 plus vitamin A, 250 µM dbcAMP, 40 ng ml–1 BDNF, 40 ng ml–1 GDNF, 200 µM ascorbic acid, 1 µg ml–1 natural mouse laminin and 2 µM arabinosylcytosine (Ara-C, Sigma-Aldrich). After 48 h, the neural selection medium was fully exchanged and after a further 48 h the medium was fully exchanged with neural maintenance medium (NMM) comprising Brainphys medium with 1× B27 plus vitamin A, 250 µM dbcAMP, 40 ng ml–1 BDNF, 40 ng ml–1 GDNF, 200 µM ascorbic acid and 1 µg ml–1 natural mouse laminin. Thereafter, every 48 h, half the medium was exchanged with NMM until day 21 post-transduction passage. All medium exchanges were performed using a Hamilton Star liquid handler set to 5 µl s–1 for aspirate and dispense as part of the NYSCF Global Stem Cell Array®. Passages were fully automated and performed on a robotic cluster comprising a Thermo Fisher Scientific C24 Cytomat incubator, a Hamilton Star liquid handler, an Agilent microplate centrifuge, a Precise Automation PreciseFlex 400 Sample Handler and a PerkinElmer Opera Phenix. At harvest, medium was removed using the Bluewasher (BlueCatBio) and cells were lysed for 5 min using RLT plus buffer (Qiagen), snap frozen on dry ice and stored at −80 °C. A replicate plate was fixed for immunofluorescence analysis by adding 32% paraformaldehyde (Electron Microscopy Sciences) directly to medium to a final concentration of 4% and incubated at room temp for 15 min. Cells were washed three times with HBSS (Thermo Fisher Scientific), stained overnight with rabbit anti-GABA 1:1000 (Sigma, catalog no. A2052), mouse anti-Nestin 1:3,000 (Millipore, catalog no. 09-0024) and chicken anti-MAP2 1:3,000 (Abcam, catalog no. 09-0006) in 5% normal goat serum (Jackson ImmunoResearch) in 0.1% Triton X-100 (Thermo Fisher Scientific) in HBSS. Primary antibodies were counterstained with goat anti-rabbit Alexa Fluor 488, goat anti-mouse Alexa Fluor 555, goat anti-chicken Alexa Fluor 647 and 10 µg ml–1 Hoechst for 1 h at room temp. Cells were washed three times with HBSS. 10 fields (×40 water objective) each with 8 planes (2µm apart) were imaged per well (one well per condition per line) using the PerkinElmer Opera Phenix microscope in confocal mode with 2x image binning (**S. Figure 4**). Cells are developmentally similar to previous hiPSC studies (**S. Figure 5A-C**).

### Glucocorticoid treatment

HCort treatment medium was prepared by first dissolving HCort (Sigma-Aldrich, catalog no. H0888) in ethanol to make a 2.8 mM stock. HCort ethanol stock was then diluted to 0.2 mM in HBSS. Ethanol was equalized to 15 µM in control and all treatment media. The final treatment medium was prepared by diluting HCort stocks into NMM, before applying to cells by fully exchanging medium. Neurons were treated with HCort for 24 h (baseline, 100 nM, 1,000 nM). HCort treatment did not impact cell number of maturity (**S. Figure 5D**).

### Genotype preprocessing

Genotype imputation was performed using the Michigan Imputation Server using the 1000 Human Genomes Project Phase 3 reference dataset using human genome build hg19. Genotypes were filtered using plink v1.9 to remove sex chromosomes, insertions/deletions, ambiguous genotypes, and retain common variants (MAF>5%), variants present in the majority of samples (missingness<10%), and variants meeting Hardy Weinberg equilibrium expectations (p> 1×10^-6^). Principal component analysis (PCA) was performed using plink v1.9 to determine genomic ancestry components which were used as covariates in downstream analysis (**S. Figure 1C**). All eQTL analyses used genotype dosages.

### RNA sequencing

RNA sequencing was performed as described previously^42^. Quality control was performed on paired-end raw sequencing reads and low-quality reads were filtered using FastQC. Short reads with Illumina adapters were trimmed using Scythe and sickle. Reads were mapped to the hg38/GRCh38 human reference genome with Rsubread. Feature-level quantification based on GENCODE release 25 (GRCh38.p7) annotation was run on alignedreads using featureCounts (subread version 1.5.0-p3) with a median 43.8% (IQR:37.3%-49.0%) of mapped reads assigned to genes.

Raw count data was filtered to remove low-expressed genes that did not meet the requirement of a minimum of 20 counts in at least ∼20% of samples. All expression values were converted to log_2_CPM and normalized to library size from mean-variance relationship estimates using edgeR v3.32.0 and limma v3.36.0. Normalized expression was subjected to unsupervised principal component analysis (PCA) to identify outliers that lay outside 95% confidence intervals from the grand averages. Variance explained by confounders such as age at death, RNA quality RIN score, post-mortem interval, and sex was determined using VariancePartition (**S. Figure 1D**). To detect hidden sources of variation in the expression data, surrogate variable analysis was performed for each brain tissue separately using sva v3.30.1 with the “be” method, preserving the effects of cumulative trauma. Surrogate variables were then residualized from normalized expression values. This tissue-specific, surrogate variable-residualized matrix was used for subsequent analyses.

### Power determinations

To ensure sufficient power to detect eQTLs in the post-mortem brain cohort, power was assessed based on effect size, variance, and sample size assumptions from the GTEx Project^50^. This study was powered at 80% to detect a linear association of genotype with a minor allele frequency (MAF)! 6.35% with expression (**S. Figure 1E**).

Given the lack of empirical data on eQTL detection in hiPSC-induced neurons, power calculations were based on effect size, variance, and sample size assumptions from post-mortem studies as above. However, these analyses are confounded by age, sex, diagnosis, medication use, post-mortem interval and more. Data in hiPSC-induced neurons has detected up to 10-fold larger effect sizes relative to post-mortem studies and 2-to-4-fold reductions in standard deviation (SD) per gene^51–53^. Assuming a 4-fold relative reduction in SD at a 20,000 SNPs-per-gene level, this study was sufficiently powered at 80% to detect an association of variants with a MAF ≥ 4.06% with expression (**S. Figure 1F**).

### eQTL detection

MatrixEQTL was used to detect variants regulating expression within a +/- 1MB cis window. Multiple testing correction was applied using a Bonferroni-corrected p-value threshold for the number of SNPs tested per gene. The genotypic contribution to expression (‘base-eQTLs’) was assessed using eqn. 1. An interaction term (eqn, 2) was modelled to identify how this regulatory relationship changed in the context of stress. Significant variants in this analysis were termed stress-interactive eQTLs. This method performs independent tests on each variant, and therefore overestimates the true number of variants contributing to gene expression due to linkage disequilibrium. Forward-stepwise conditional analysis was performed to identify conditionally independent signals.

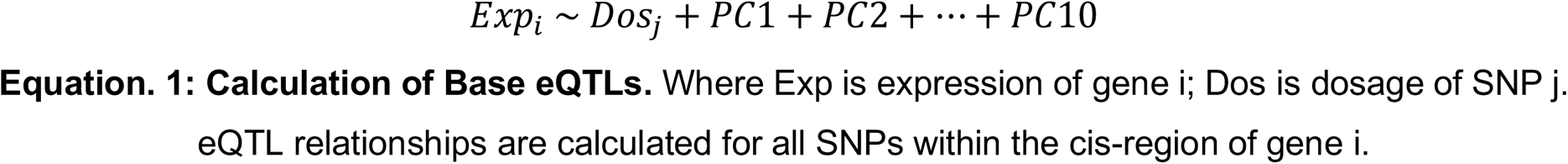

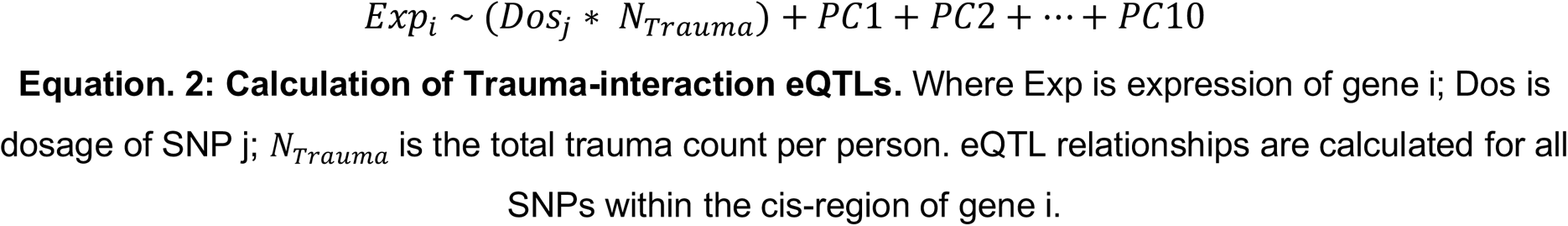

### eQTL linear model assumption testing

Interactive eQTLs are susceptible to unequal variance at extreme ends of environmental exposure and with increasing dosage of the minor allele, leading to violation of linear regression assumptions. Ordinary least squares regression (OLS) ensured the robustness of stress-interactive eQTLs. There are four assumptions of OLS: linearity of the predictor and response, independence, normality of the errors, and homoscedasticity (constant error variance). The relationships of SNP dosage and expression are assumed to be linear, as is the relationship between expression and traumatic exposure. Independence also is satisfied as the data is not time-series. The assumption of normality was statistically tested for each model using the Shapiro-Wilk test in R. The assumption of homoscedasticity was assessed using the Koenker’s studentized version of the Breusch-Pagan test, bptest() in the lmtest package, v 0.9-38. Significance in either of these tests indicates the samples come from a non-normal or heteroskedastic population, respectively. A model was deemed to meet the assumptions if the test statistic of both tests was non-significant at a p value threshold of 0.05.

### eQTL replication

Overlap between eQTLs and eGenes identified in our analysis with Bonferroni-significant associations in previous Common Mind Consortium^53^ and GTEx^50^ analyses (**S. Figure 2A**) was assessed for matching tissues using percentage overlap over SNP-gene pairs, eGenes, and using the pi1 statistic from the qvalue package (v 2.33.0).

### Forward step-wise conditional analysis

To identify conditionally independent eQTLs, step-wise conditional analysis was performed^54^. Briefly, eQTLs were first associated using MatrixEQTL. Significance was initially assessed using a Bonferroni-corrected threshold across all *cis*-eQTL tests within each gene. P values were not re-assessed at each conditional step; instead, a fixed p value threshold was used as the inclusion criteria in the stepwise model selection. For each gene with at least one *cis*-eQTL (gene ± 1 Mb) association at a 5% false discovery rate (FDR), the most significant SNP was added as a covariate in order to identify additional independent associations (considered significant if the p value achieved was less than that corresponding to the Bonferroni threshold for primary eQTL). This procedure was repeated iteratively until no further eQTL met the p value threshold criteria (**S. Figure 2B**).

### Pair-wise eQTL colocalization

Pair-wise colocalization using coloc^55^, with loci defined by each lead SNP ± 1 Mb, ddetermine whether or not eQTL architecture was conserved across conditions (brain regions, cell types, exposure to HCort) (**S. Figure 2C**). To determine whether shared significant eGenes had equivalent underlying genetic regulation under multiple conditions (ie: across different brain regions, in the brain vs in hiPSC-derived neurons, or in hiPSC-derived neurons treated with varying amounts of HCort), genes with a PPH4>0.8 were selected. To determine eGenes that were significant in multiple conditions but due to different underlying genetic regulation, genes with a PPH3>0.8 were selected (**S. Figure 2D**).

### Gene set enrichment

Gene set enrichment of GO biological processes, cellular components, and molecular function, and genome-wide association studies (GWAS) catalog reported genes, was performed using FUMA^56^. Where relevant, stress-interactive eGene enrichments were always performed against base eGenes as a background. If not specified, the background gene list used was expressed genes in each relevant post-mortem brain region as defined by genes that met a minimum of 20 counts in ∼20% of samples.

### Cell type-specific expression imputation

To determine cell type proportions of the four post-mortem brain regions included in our analysis, log-2-normalized TPM expression matrices were input to CIBERSORTx^57^ alongside PsychENCODE single cell reference gene signatures^58–60^. Cell type proportions derived from CIBERSORTx were input to bMIND^61^ alongside the bulk RNA log-2-normalized expression matrix to derive imputed cell type specific expression. To detect hidden sources of variation in imputed expression data, sva^62^ was performed for each cell type-imputed expression matrix. This tissue-specific, surrogate variable-residualized matrix was used for subsequent analyses.

### Motif enrichment

Lead SNPs from both base and stress-interactive conditional analysis and any SNPs in high LD>0.8 were used to query Haploreg^63^ to determine motifs disrupted by each SNP. Motifs more often disrupted by stress-interactive eQTLs were determined using a binomial test.

### Transcription factor validation

Validated control hiPSC-derived NPCs for RNAi were selected from a previously reported case/control hiPSC cohort of childhood onset SCZ (COS): NSB553-S1-1 (male, European ancestry), NSB2607-1-4 (male, European ancestry). hiPSC-NPCs were generated and validated as previously described (ref) and cultured in hNPC media (DMEM/F12 (Life Technologies #10565), 1x N2 (Life Technologies #17502-048), 1x B27-RA (Life Technologies #12587-010), 1x Antibiotic-Antimycotic, 20 ng/ml FGF2 (Life Technologies)) on Matrigel (Corning, #354230). hiPSC-NPCs at full confluence (1-1.5×10^7^ cells / well of a 6-well plate) were dissociated with Accutase (Innovative Cell Technologies) for 5 mins, spun down (5 mins X 1000g), resuspended, and seeded onto Geltrex-coated plates at 3-5×10^6^ cells / well. Media was replaced every two-to-three days for up to seven days until the next split.

At day -1, NPCs were transduced with rtTA (Addgene 20342) and *NGN2* (Addgene 99378) lentiviruses. Medium was switched to non-viral medium four hours post infection. At day 0 (D0), 1 µg/ml dox was added to induce *NGN2*-expression. At D1, transduced hiPSC-NPCs were treated with antibiotics to select for lentiviral integration (300 ng/ml puromycin for dCas9-effectors-Puro, 1 mg/ml G-418 for NGN2-Neo). At D3, NPC medium was switched to neuronal medium (Brainphys (Stemcell Technologies, #05790), 1x N2 (Life Technologies #17502-048), 1x B27-RA (Life Technologies #12587-010), 1 µg/ml Natural Mouse Laminin (Life Technologies), 20 ng/ml BDNF (Peprotech #450-02), 20 ng/ml GDNF (Peprotech #450-10), 500 µg/ml Dibutyryl cyclic-AMP (Sigma #D0627), 200 nM L-ascorbic acid (Sigma #A0278)) including 1 µg/ml Dox. 50% of the medium was replaced with fresh neuronal medium once every 2-3 days.

On day 5, young neurons were replated onto geltrex-coated 12-well plates at 1.2×10^^6^ cells / well. Cells were dissociated with Accutase (Innovative Cell Technologies) for 5-10 min, washed with DMEM, gently resuspended, counted and centrifuged at 1,000xg for 5 min. The pellet was resuspended at a concentration of 1.2×10^6^ cells/mL in neuron media [Brainphys (StemCell Technologies #05790), 1xN2 (ThermoFisher #17502-048), 1xB27-RA (ThermoFisher #12587-010), 1 mg/ml Natural Mouse Laminin (ThermoFisher #23017015), 20 ng/mL BDNF (Peprotech #450-02), 20 ng/mL GDNF (Peprotech #450-10), 500 mg/mL Dibutyryl cyclic-AMP (Sigma #D0627), 200 nM L-ascorbic acid (Sigma #A0278)] with doxycycline. Doxycycline was fully withdrawn from the neuronal media at day 7.

At D13, iGLUTs were treated with 200 nM Ara-C to reduce the proliferation of residual non-neuronal cells in the culture, followed by half medium changes. At D17, Ara-C was completely withdrawn by full medium change while adding media containing pooled shRNA vectors, either targeting *YY1*, *MYC*, or scramble controls. Medium was switched to non-viral medium four hours post infection. At D18, transduced iGLUTs were treated with corresponding antibiotics to the shRNA lentiviruses (300 ng/ml puromycin). 24 hours prior to harvest (D20), iGLUTs were treated with 1 μM hCort or ethanol vehicle control. Neurons were harvested for RNA extraction and bulk RNA-seq at D21. Samples from 3 wells per condition per donor were harvested in total (6 samples per condition total).

### Neuronal pooled CRISPRi screen

128 putative regulatory regions were prioritized corresponding to 65 eGenes based on minimizing distance to transcription start site, presence in glutamatergic neuron open chromatin, and in GR binding sites, obtained from ReMap2022. gRNA design was conducted using Benchling and CRISPR-ERA. gRNAs were selected based on their lack of predicted off targets and E scores (**Table S3**). Generation of the CRISPRi guide library and lentiviral packaging were outsourced to GenScript. Briefly, gRNA DNA oligos were synthesized and cloned into the lentiGuide-Hygro-mTagBFP2 plasmid which was confirmed by next generation sequencing. Plasmids were packaged into lentivirus at a titer of 1.34×10^8^.

For pooled analysis, validated control (from donors 2607 and 553, as above) hiPSC-derived NPCs expressing dCas9-KRAB for CRISPRi were transduced with rtTA (Addgene 20342) and *NGN2* (Addgene 99378) lentiviruses and grown as described above. At D17, pools of mixed gRNA vectors (Addgene 99374), either targeting relevant genes or scramble controls, were added to the neuronal media. Medium was switched to non-viral medium four hours post infection. At D18, transduced iGLUTs were treated with corresponding antibiotics to the gRNA lentiviruses (1 mg/ml HygroB for lentiguide-Hygro/lentiguide-Hygro-mTagBFP2). 24 hours prior to harvest (D20), iGluts were treated with 1 μM hCort or an ethanol vehicle as a control. At D21, iGLUT neurons were dissociated to single cell suspensions with papain, antibody-hashed, and bar-coded single cell cDNA generated using 10X Genomics Chromium in order to perform ECCITE-seq. 20,000 cells were sequenced per sample, with four samples in total submitted for single cell sequencing (one per condition per donor).

<u>E</u>xpanded <u>C</u>RISPR-compatible <u>CITE-seq</u> (ECCITE-seq) combines Cellular Indexing of Transcriptomes and Epitopes by sequencing (CITE-seq) and Cell Hashing for multiplexing and doublet detection with direct detection of sgRNAs to enable single cell CRISPR screens with multi-modal single cell readout. By capturing pol III-expressed guide RNAs directly, this approach overcomes limitations of other single-cell CRISPR methods, which detect guide sequences by a proxy transcript, resulting in barcode switching and lower capture rates.

### Analysis of single-cell CRISPRi screen

Single cell sequencing reads were mapped to the GRCh38 reference genome using Cellranger 3.1.0. Kallisto^64^ was used to generate HTO counts and index GDO libraries. Seurat (v.2.3.0) was used for QC, normalization, cell clustering, HTO/GDO demultiplexing, and DEG analysis. Each sequencing lane was initially processed separately. Cells with RNA UMI feature counts were filtered (200 < nFeature_RNA < 8000) and the percentage of all the counts belonging to the mitochondrial, ribosomal, and hemoglobin genes calculated using Seurat::PercentageFeatureSet. Hashtag and guide-tag raw counts were normalized using centered log ratio transformation. For demultiplexing based on hashtag, Seurat::HTODemux function was used; and for guidetag counts Seurat::MULTIseqDemux function within the Seurat package was performed with additional MULTIseq semi-supervised negative-cell reclassification. To remove variation related to cell-cycle phase of individual cells, cell cycle scores were assigned using Seurat::CellCycleScoring. RNA UMI count data was then normalized, log-transformed and the percent mitochondrial, hemoglobulin, and ribosomal genes, cell cycle scores (Phase) regressed out using Seurat::SCTransform (**S. Figure 6D**). Lanes were then integrated using Harmony 1.0^65^. To ensure that cells assigned to a guide-tag identity class demonstrated successful perturbation of the target gene, ‘weightednearest neighbor’ (WNN) analysis was performed to assign clusters based on both guide-tag identity class and gene expression. To identify successfully perturbed cells, pseudobulking was performed using donor as a grouping factor and calculated differential expression of each perturbed gene using edgeR v3.32.0 and limma v3.36.0.

### Transcriptomic imputation

Elastic net regression with ten-fold cross validation was used to generate transcriptomic imputation models for each brain region of interest, using the glmnet package^66^. An 11^th^ hold-out fold was used for testing. All cross-validation folds were balanced for diagnoses, ancestry, and other clinical variables. Two models were generated for each brain region; (1) a “base” model, where only SNP dosages were included in the regression analysis, and (2) a “stress-interactive” model, where SNP dosages, total traumatic event count, and the interaction between SNP dosages and traumatic exposures (SNP * trauma) for all SNPs with a nominally significant interaction term (p<0.05) in the eQTL analysis were included in the regression analysis. For both models, only SNPs within the cis-region (±1 Mb) of each gene were included in the regression analysis. Accuracy of prediction was first estimated by comparing predicted expression to measured expression, across all ten cross-validation folds; this correlation was termed cross-validation *R*^2^ or*Rcv*^2^. Next, accuracy was measured by predicting expression of the 11^th^ hold-out fold, and comparing predicted expression to measured expression. This correlation was termed within-sample validation *R*^2^. Genes with *Rcv*^2^ > 0.01 and *P* < 0.05) were included in our final predictor database.

### Application of transcriptomic imputation models to external biobanks

Elastic net models were applied to the Bio*Me*™ biobank and the UK Biobank. The Bio*Me*™ biobank consists of 28,250 individuals with matched genotype and electronic health records. The Bio*Me*™ cohort is racially diverse (self-reported race: Hispanic American, 35%; European American, 34%; African American, 25%; Other 6%), with an average age of 59.84 (SE=17.85). As traumas are known to be unequally distributed across the population by race, expression was imputed separately by race and meta-analyzed together. Traumatic experiences in this cohort were defined through the EHR as described previously^67^.

UK Biobank is a national resource sampled from 22 assessment centers across the UK. Traumatic experiences were defined from structured interview completion of the mental health questionnaire, from positive answers to fields 20488, 20490, 20523, 20521, 20524, 20531, 20529, 20526, 20530, 20528, 20527, 20487, and negative answers to 20491 and 20489. 157,322 individuals who completed this survey were included in this analysis.

Genotype preprocessing was conducted as previously described here. Genotype dosages were scaled, and weights from each model were applied to each SNP, and summed to generate imputed expression for each gene. Next, expression was assessed for 12 binary case-control traits based on presence of ICD codes (PTSD: F43, MDD: F32, F33, ANX: F40, F41, SCZ: F20, ASD: F84, ADHD: F90, BP: F31, AN: F50, OCD: F42, AD: G30, AUD: F10). 10 genotype-derived PCs, age, and sex were used as covariates.

## RESULTS

### Genetic regulation of gene expression is modified by traumatic stress

Baseline and trauma-interactive genetic regulatory relationships were examined across four stress disorder-related post-mortem brain regions (n=304 donors): the dorsolateral prefrontal cortex (DLPFC), the dorsal anterior cingulate cortex (dACC), the basolateral amygdala (BLA), and the medial amygdala (MeA). Across all four brain regions, 655,607 significant cis-eQTLs were identified, including 7,964 unique eGenes (**Figure 1C**). We term these trauma-independent associations **base eQTLs** and their target genes **base eGenes.** Base eQTLs significantly overlapped previously reported post-mortem brain eQTLs; 23.03% overlap with GTEx-DLPFC eQTLs (36.94% of eGenes; pi1 statistic: 0.85), and 18.86% overlap with GTEx-ACC eQTLs (31.09% of eGenes, pi1 statistic: 0.78). No BLA and MeA samples are available in GTEx; however, 15.19% of BLA eQTLs (24.76% of eGenes, pi1 statistic: 0.77) and 15.50% of MeA eQTLs (25.09% of eGenes, pi1 statistic: 0.78) overlap with bulk GTEx amygdala measures (**S. Figure 2A**).

**Figure 1:**
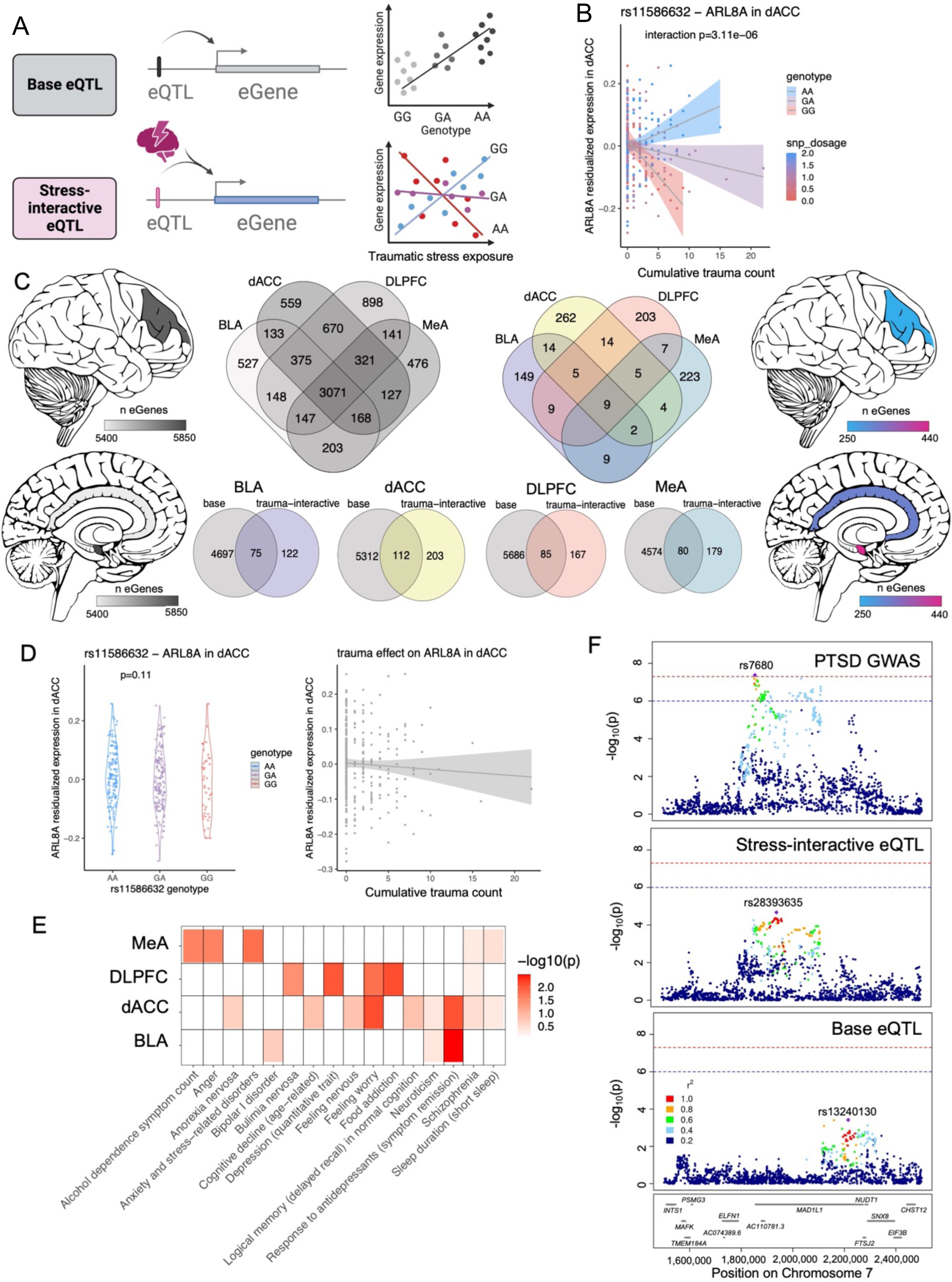
Genetic regulation of expression is modified by traumatic stress. A) Base eQTLs linearly associate genotype with expression of a nearby gene. Stress-interactive eQTLs incorporate a quantitative measure of stress exposure in a linear interaction term with genotype to measure the modification of the eQTL relationship in the context of traumatic stress. B) rs11586632 is a stress-interactive eQTL for the ARL8A eGene in the dACC, where ARL8A expression is differentially impacted by increasing stress burden in individuals with different genotypes. C) Cerebroviz images representing the number of base eGenes (left) and stress-interactive eGenes (right) detected across post-mortem brain regions. Venn diagrams show overlap of base eGenes (left) and stress-interactive eGenes (right) across the post-mortem brain. Venn diagrams (below) show overlap of base eGenes and stress-interactive eGenes across post-mortem brain regions. D) (left) ARL8A expression is not significantly associated with rs11586632 genotype, demonstrating it is not a base eQTL. (right) ARL8A expression is not significantly associated with increasing traumatic stress burden, indicating it is not a stress-induced differentially expressed gene. E) Gene set enrichment of stress-interactive eGenes across post-mortem brain regions in gene sets of neuropsychiatric traits from the GWAS catalog. F) Locuszoom of the *NUDT1* locus showing SNP associations with PTSD from the PTSD GWAS (top), stress-interactive SNP associations with *NUDT1* gene expression (middle), and base (non-stress associated) SNP associations with *NUDT1* gene expression (bottom). Red dotted line indicates genome-wide significance.

Most base eGenes were shared across brain regions (69.1%), albeit with more overlap between cortical regions than amygdala regions. Of overlapping eGenes, between 458 and 712 eGenes exhibited conditionally independent genetic regulation among tissue pairs, indicating divergent regulatory mechanisms despite shared eGenes (**S. Figure 2C, D**).

To assess how baseline regulatory relationships differ with degree of exposure to traumatic stress, the interaction effect between genotype dosage and cumulative trauma count on gene expression was tested (**Figure 1A**). We term eQTLs with a significant interaction effect **stress-interactive eQTLs** and their target genes **stress-interactive eGenes.** 8,557 **stress-interactive eQTLs** were identified across the post-mortem brain, corresponding to 915 unique **stress-interactive eGenes**. Significant stress-interactive eQTLs were not simply genes with differential expression due to trauma; 16%, 6%, 11%, and 16% of stress-interactive eGenes were nominally significant DEGs with traumatic exposure in the BLA, dACC, DLPFC, and MeA, respectively. Likewise, stress-interactive eQTLs were not always significant base eQTLs; 38%, 36%, 34%, and 31% of stress-interactive eGenes were base eGenes in BLA, dACC, DLPFC, and MeA, respectively (e.g., rs11586632, **Figure 1B**).

### Genes with underlying stress-interactive genetic regulation show region-specificity in enrichment for neuropsychiatric traits

Stress-interactive eGenes, relative to base eGenes, were less likely to be shared between brain regions; only 8.5% of stress-interactive eGenes had a significant stress-interactive eQTL in multiple brain regions, compared to 69.1% of base eGenes (**Figure 1C**). These differences were not driven by variance in cell type proportion between brain tissues (**S. Figure 3A**), as after adding region-specific cell type proportions as a covariate, 87.3 - 90.1% of stress-interactive eGenes and 97.4 - 98.0% of base eGenes remained significant, suggesting that these eGenes were not largely explained by cell type proportion (**S. Figure 3B**). Overall, stress-interactive eGenes were less likely than base eGenes to be shared between brain regions, suggesting region-specificity in genetically regulated encoding of traumatic experiences.

Stress-interactive eGenes, compared to base eGenes, were differentially enriched in gene sets associated with neuropsychiatric traits, including ‘feeling worry’, in cortical regions (p=1.68×10^-2^ in DLPFC, p=8.04×10^-3^ in dACC), and ‘anxiety and stress-related disorders’ in the MeA (p= 1.72×10^-2^) (**Figure 1E**). Moreover, stress-interactive variants differentially colocalized with PTSD GWAS loci; for example, in the BLA, stress-interactive eQTLs colocalized with PTSD GWAS variants around the *NUDT1* locus (PPH4=0.838), while baseline genetic regulation did not (PPH4=0.023) (**Figure 1F**), suggesting that PTSD risk at this locus is reflective of variants modified by traumatic stress.

### Stress-interactive variants lie in stress-related transcription factor binding domains

To mechanistically examine how stress-interactive eQTLs confer genotype-dependent regulation of stress encoding (**Figure 2A**), unbiased enrichment of stress-interactive eQTLs compared to base eQTLs in known transcription factor binding sites was performed. 30, 52, 54, and 83 out of 1188 motifs^68^ were significantly enriched (binomial p<0.05) for stress-interactive eQTLs relative to base eQTLs in the BLA, dACC, DLPFC, and MeA, respectively (**Table S3**). Significantly enriched transcription factors included effectors of glucocorticoid-mediated stress^69,70^ such as NFkB (BLA p=1.37×10^-4^, DLPFC p=4.3×10^-3^) and GR (BLA p=1.31×10^-3^, DLPFC p=4.5×10^-2^), and also included transcription factors mediating PTSD diagnosis-dependent transcriptional hypersensitivity to glucocorticoids in iGLUTs^31^: YY1 (p=3.3×10^-2^) and MYC (p=4.75×10^-2^) in the DLPFC (**Figure 2F**). Together, this suggests that stress-interactive eQTLs exert genotype-dependent response to traumatic stress due to their position in stress-relevant transcription factor binding sites.

**Figure 2:**
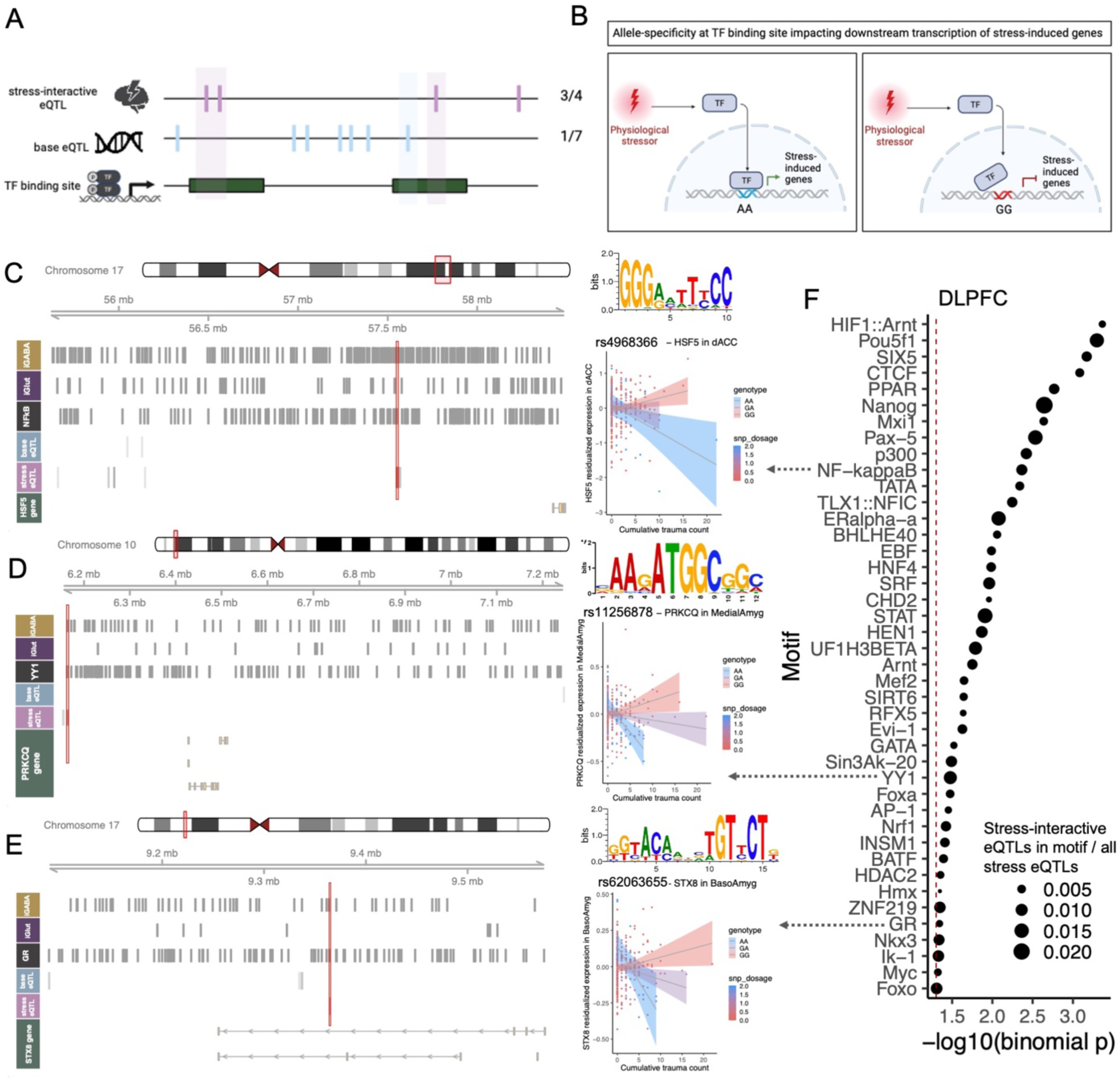
Unbiased discovery of novel transcription factors identifies mediators of stress response. A) Schematic showing enrichment of stress-interactive eQTLs in transcription factor binding sites was determined by frequency of stress-interactive eQTLs in that transcription factor’s binding sites against frequency of base eQTLs in that transcription factor’s binding sites. B) Proposed mechanism for genotype-dependent disruption of transcription factor binding sites, leading to differential regulation of stress-induced gene expression. C) rs4968366 is the lead stress-interactive eQTL for the HSF5 eGene in the dACC. rs4968366 falls in an NFkB binding motif in iGABA open chromatin. Base eQTLs for this eGene do not fall in NFkB binding sites. D) rs11256878 is the lead stress-interactive eQTL for the PRKCQ eGene in the MeA. rs11256878 falls in a YY1 binding motif in iGABA open chromatin. Base eQTLs for this eGene do not fall in YY1 binding sites. E) rs62063655 is the lead stress-interactive eQTL for the STX8 eGene in the BLA. rs62063655 falls in a GR binding motif in iGABA open chromatin. Base eQTLs for this eGene do not fall in GR binding sites. F) Transcription factors enriched for stress-interactive eQTLs within binding motifs compared to base eQTLs in the DLPFC. Size of point indicates the ratio of stress-interactive eQTLs across the genome lying in the indicated motif compared to the total number of stress-interactive eQTLs.

### Genetic regulation of glucocorticoid-mediated stress in vitro replicates molecular mechanisms of stress encoding in the post-mortem brain

As post-mortem brain findings may be confounded by diverse lifetime experiences and heterogeneity of traumatic exposures, our findings were validated using an *in vitro* model of stress exposure, whereby donor-matched hiPSC-derived neurons from PTSD cases and controls (n=39) are treated with HCort^31^ (0 nM, 100 nM, and 1000 nM) (**Figure 3A**). We focused on glutamatergic and GABAergic cell types, as they are highly implicated in PTSD^71–73^; these cell types were generated via transient overexpression of *NGN2* in hiPSCs to yield iGLUT neurons (typically >95% yield and functionally mature by day 21-28^49,51,74–77^) or transient overexpression of *ASCL1* and *DLX2* to yield iGABA neurons (typically >80% yield and functionally mature by day 35-42^49,77^) (**S. Figure 4**).

**Figure 3:**
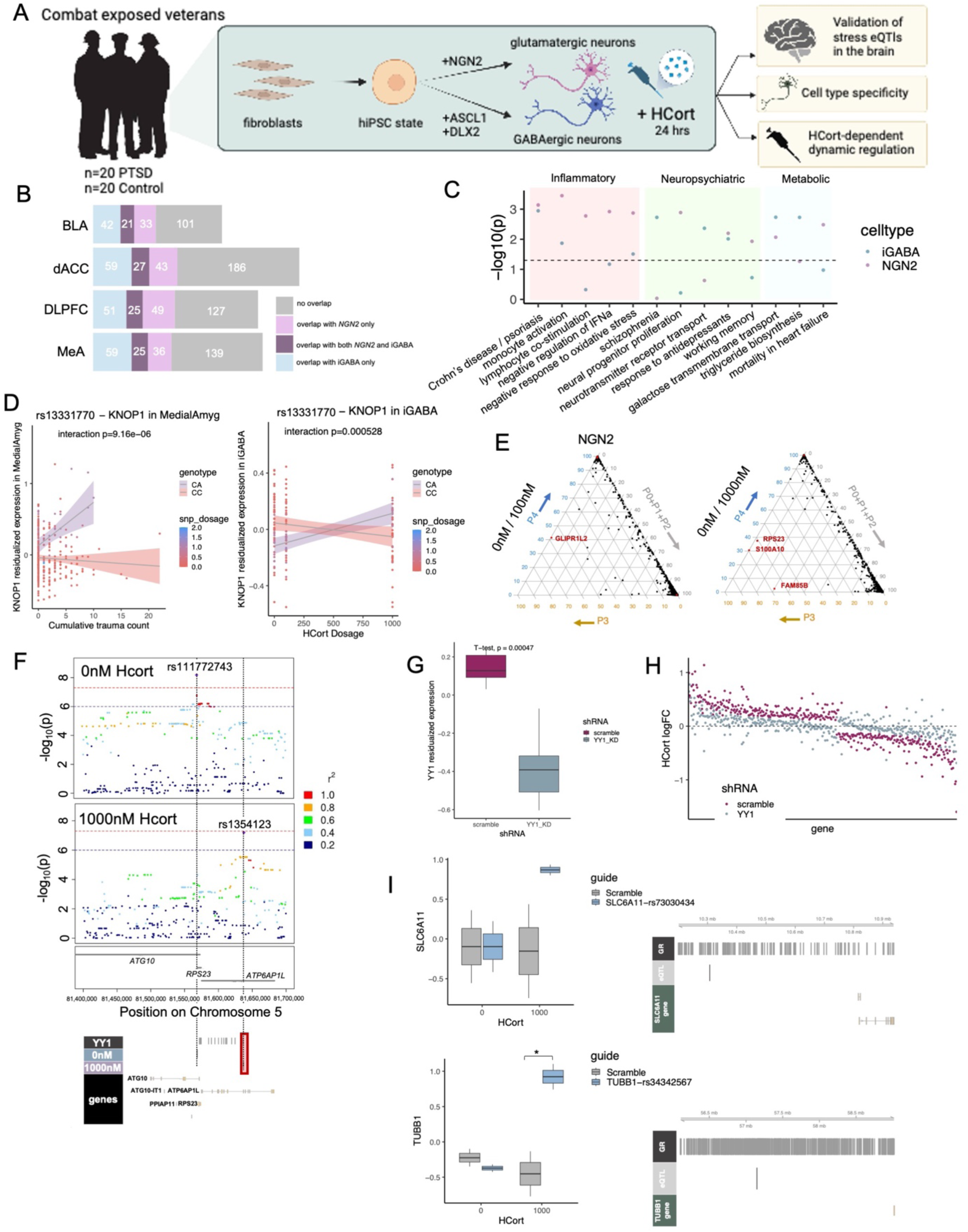
hiPSC-derived neurons validate genetic regulation of glucocorticoid-mediated stress. A) Schematic showing collection and reprogramming of PTSD case and control donor fibroblasts through the hiPSC state and differentiation into glutamatergic and GABAergic neurons, treated with HCort for 24 hours. B) Overlap of glucocorticoid dosage-interactive eGenes in hiPSC (blue: overlap with iGABA, pink: overlap with iGLUT, purple: overlap with both) with stress-interactive eGenes across the four post-mortem brain tissues. C) Enrichment of unique iGABA (blue) and iGLUT (pink) stress-interactive eGenes in inflammatory, neuropsychiatric, and metabolic pathways. Black dotted line indicates significant enrichment. D) rs13331770 is a stress-interactive eQTL regulating the KNOP1 gene in the MeA with traumatic stress (left), and in GABAergic neurons with glucocorticoid-induced stress (right). E) Ternary plots showing pair-wise colocalization of glutamatergic neurons treated with 100 nM HCort (left) and 1000 nM HCort (right) compared to baseline. P0/P1/P2 indicates no eQTL association at either dose or association only at a single dose. P3 indicates eQTL associations at both doses but at separate causal variants. P4 indicates eQTL associations at both doses with a shared causal variant. Genes with P3>0.5 are highlighted in red. F) Locuszoom of the *RPS23* locus showing SNP associations with *RPS23* gene expression at baseline (top), and with exposure to 1000 nM HCort (bottom). Red dotted line indicates genome-wide significance. The lead SNP at the *RPS23* locus at baseline (rs111772743, blue track) and with 1000 nM HCort (rs1354123, purple track) indicating that only the lead SNP with HCort exposure (rs1354123) lies in a YY1 binding motif. G) YY1 expression is reduced with shRNA-mediated knockdown compared to scramble. H) The log2FC of HCort-stimulation in HCort-responsive genes in the scramble condition (purple) and with YY1 shRNA-mediated knockdown (blue). I) (left) Expression of eGenes treated with scramble (gray) or CRISPRi perturbation (blue) of putative stress-interactive regulatory regions at baseline or with HCort exposure. Targeted regions are located in GR binding sites (right).

We define **hCort-interactive eGenes** as those with neuronal context-specific regulation *in vitro*. 41-50% of trauma-interactive eGenes in the post-mortem brain were also identified as hCort-interactive eGenes *in vitro* (**Figure 3B**) (e.g., rs13331770, MeA p=9.16e-6, iGABA p=5.28e-4, **Figure 3D**); the majority of these overlapping eGenes are stress-interactive in either iGABA or iGLUT cells. (**Figure 3C**). These eGenes show enrichments across inflammatory, neuropsychiatric, and metabolic pathways (**Figure 3D**) that are both shared (e.g., response to antidepressants, iGLUT p=6.32×10^-3^, iGABA p=9.72×10^-3^) and distinct (e.g., neurotransmitter receptor transport, iGLUT p=0.235, iGABA p=4.31×10^-3^) across cell types (**Figure 3D**).

Many hCort-interactive eGenes (*RPS23, CTNNB1, GLIPR1L2, S100A10,* and *FAM85B*) showed distinct eQTL regulatory architecture at baseline compared to following glucocorticoid exposure (PPH3>0.5; **Figure 3E, S. Figure 6**). For example, two distinct SNPs associated with *RPS23* expression; the lead SNP (rs1354123) regulating *RPS23* expression in the presence of HCort-mediated stress lies in a YY1 transcription factor binding site, while the lead SNP (rs111772743) at baseline does not (**Figure 3F**). We previously identified YY1 targets as enriched for glucocorticoid-hypersensitivity in PTSD cases^31^, and here identified YY1 binding sites as enriched for stress-eQTLs compared to base-eQTLs in the post-mortem DLPFC (**Figure 2F**); consistent with this, shRNA-mediated knockdown of YY1 (**Figure 3G**) reduced the sensitivity of hCort-responsive genes to hCort (**Figure 2H**), suggesting that YY1 mediates glucocorticoid-dependent transcriptomic responses. This was likewise true for MYC, which we previously implicated in PTSD-mediated hyper-responsivity^31^ (**S. Figure 6C**).

Stress-interactive eGenes shared between hiPSC-derived neurons and the post-mortem brain (by functional annotation of lead eQTLs in known GR binding sites and iGLUT open chromatin^78^) were repressed in a single cell CRISPR-inhibition screen^79^. HCort-dependent activity of putative regulatory elements, such as the rs34342567 SNP, conferred positive regulatory activity of the *TUBB1* gene only with hCort exposure (interaction p=1.8×10^-2^), suggesting this site mediates hCort hyper-responsivity (**Figure 3I**).

### Glial and endothelial cells are enriched for genotype-dependent stress response

Given the cell type-specificity of transcriptomic response to glucocorticoid-mediated stress in hiPSC-derived neurons, we next considered whether bulk tissue eQTLs were driven by cell type-specific regulation. To do this, cell-type-specific gene expression in post-mortem tissues was deconvoluted from bulk expression, then base and trauma-interactive eQTL analyses were repeated across seven deconvoluted brain cell types: astrocytes, endothelial cells, excitatory neurons, inhibitory neurons, microglia, oligodendrocytes, and other neurons (not strictly inhibitory or excitatory) **(Figure 4A, S. Figure 3C)**. Many imputed excitatory and inhibitory neuronal stress-interactive eGenes from the post-mortem brain were replicated hiPSC-derived HCort-interactive eGenes; inhibitory neurons demonstrated greater replication (44.7-52.9% overlap) compared to excitatory neurons (26.8-34.6% overlap) (**Figure 4B)**. Stress-interactive eGenes identified in both excitatory neurons and iGLUT neurons enriched in regulatory pathways (i.e. regulation of protein folding, p=5.22×10^-4^) and oxidative stress pathways (response to oxidative stress, p=1.39×10^-3^), while iGABA enrichments included hippocampal and limbic system development (p=3.55×10^-4^ and p=7.39×10^-4^, respectively), brain regions associated with fear encoding^13,80^, in addition to stress signaling (p=1.71×10^-3^) (**Figure 4C**).

**Figure 4:**
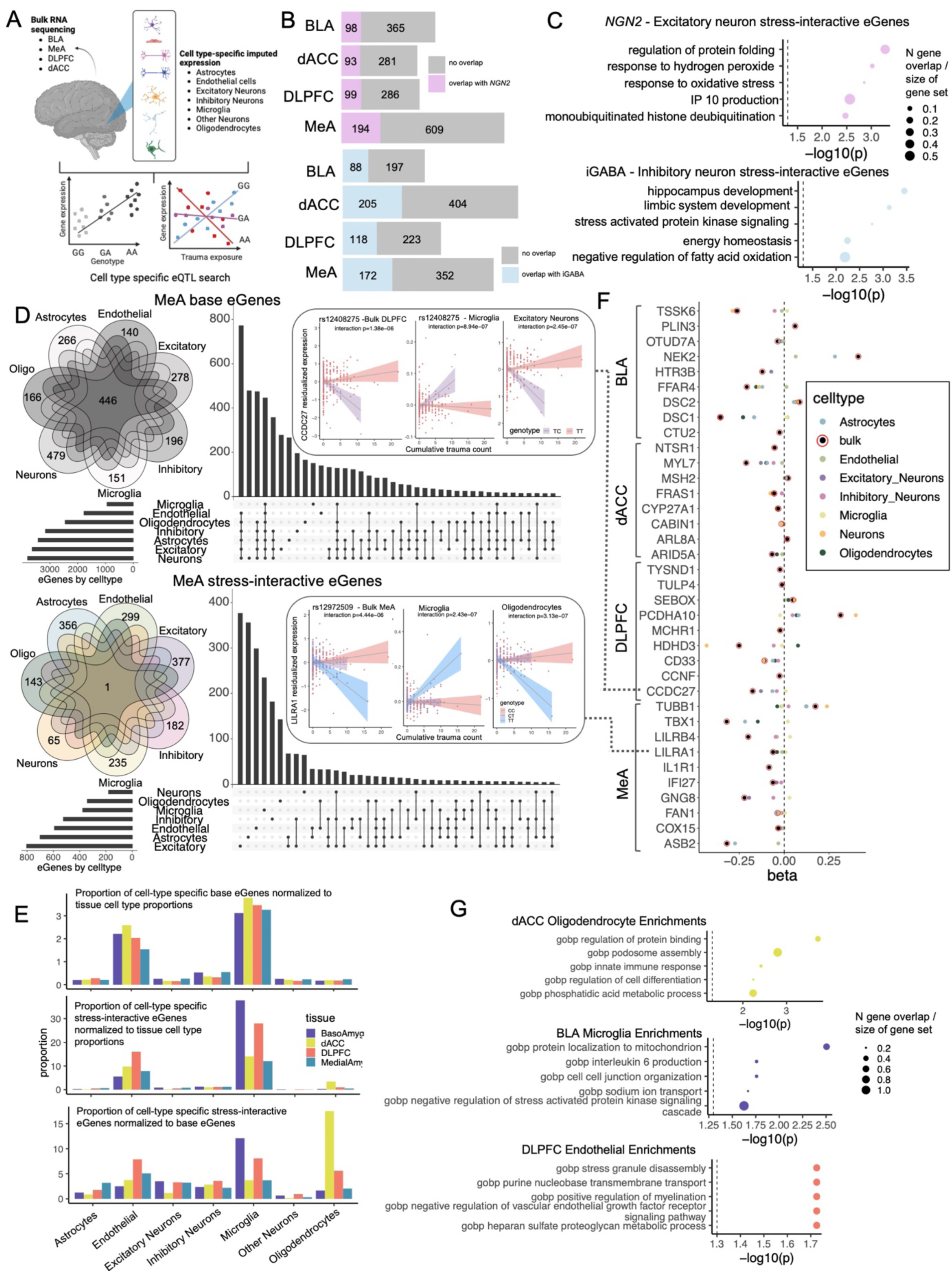
Novel cell types mediate region-specificity in genotype-dependent stress response. A) Schematic showing bulk sequencing data was deconvoluted into 7 cell type-specific expression matrices, and base- and stress-interactive eQTLs were determined for each cell type in each brain region. B) Overlap of imputed excitatory neuron stress-interactive eGenes in the post-mortem brain with hiPSC-derived iGLUT stress-interactive eGenes (top) and imputed inhibitory neuron stress-interactive eGenes with iGABA neuron stress-interactive eGenes (bottom). C) Gene ontology enrichments of overlapping stress-interactive eGenes between post-mortem brain imputed cell types and hiPSC-derived neurons for iGLUT neurons and brain excitatory neurons (top) and iGABA neurons and brain inhibitory neurons (bottom). D) Base (top) and stress-interactive (bottom) eGene overlap across imputed cell types from the MeA. E) Effect sizes for eQTLs in bulk tissue and in any cell types for which the eQTL is significant in a subset of genes across the four post-mortem brain regions, with inserts for rs12408275, a stress-interactive eQTL which has opposite effect sizes on the CCDC27 eGene in microglia compared to excitatory neurons, and rs12972509, a stress-interactive eQTL which has opposite effect sizes on the LILRA1 eGene in microglia compared to oligodendrocytes. F) Proportions of cell type-specific eGenes compared to all cell type eGenes normalized to their respective cell type proportions in the four post-mortem brain tissues for base eGenes (top), stress-interactive eGenes (middle), and in stress-interactive eGenes compared to base eGenes (bottom). G) Gene ontology enrichments of stress-interactive eGenes in oligodendrocyte in the dACC (top), microglia in the BLA (middle), and endothelial cells in the DLPFC (bottom). Dotted line indicates significance.

Cell type-deconvoluted base eQTLs largely replicate reported single-cell eQTLs, with 35% (microglial) to 62% (neuronal) overlap with previous reports^81^ (**S. Figure 3D**). Across brain regions, 69.9%-77.4% of base eGenes were shared between more than one cell type, with 10.5%-11.9% of eGenes shared between all seven cell types, respectively. In contrast, in the MeA, only a single stress-interactive eGene was shared between all cell types (**Figure 4D, S. Figure 7**).

When cell types enriched for eQTL activity were considered relative to cell type proportion, the role of endothelial cells and microglia in both base and stress-interactive eGenes across all four brain regions was highlighted (i.e. microglia DLPFC base eGene enrichment p=2.987e-34). Inhibitory neurons were enriched only in stress-interactive eGenes in the MeA and BLA (i.e. inhibitory neurons BLA trauma eGene enrichment p=3.02×10^-3^) and oligodendrocytes were enriched only in stress-interactive eGenes in the dACC and DLPFC (i.e. oligodendrocytes dACC trauma eGene enrichment p<2.2e-16) (**Figure 4E**). When comparing stress-interactive eGenes to base eGenes, stress-eGenes were particularly enriched in oligodendrocytes in the dACC (enrichment ratio: 17.50), microglia in the BLA (enrichment ratio: 12.09), and endothelial cells in the dACC (enrichment ratio: 3.753) (**Figure 4E**). Stress-interactive eGenes in these regions enriched for key processes involved in myelination^82^ (i.e. phosphatidic acid metabolism in dACC oligodendrocytes, p=5.84×10^-3^, positive regulation of myelination in DLPFC endothelial cells, p= 1.88×10^-2^) and blood brain barrier permeability^83,84^ (i.e. IL6 production in BLA microglia, p=1.74×10^-2^, heparan sulfate metabolism in DLPFC endothelial cells, p=1.88×10^-2^) (**Figure 4G**), suggesting that these cell types and processes may underlie regional specificity of genotype-dependent stress encoding.

To assess whether stress-interactive eQTLs in bulk post-mortem brain tissue might represent a linear aggregation of cell type-specific stress response, we determined the effect sizes for the most significant stress-interactive eQTL for each eGene, in each cell type for which the SNP was a significant stress-interactive eQTL. For 47.5% of stress-interactive eGenes, the top eQTL in bulk tissue was more significant than in each deconvoluted cell type (e.g. *DSC1*, **Figure 4F**). In such cases, the effect size of the bulk eQTL often represented a linear combination of several cell types (e.g. *NEK2* for astrocytes and endothelial cells). In the remainder of cases, there was evidence of cell-type-specific drivers (e.g. *CCDC27* showed a significant stress-interactive effect in the same direction as the bulk DLPFC in excitatory neurons, but opposing direction in microglia, **Figure 4F**). Notably, the cell type with the most significant trauma-interactive eQTL was largely not the cell type with the largest cell fraction.

### Genotype-dependent stress encoding implicates novel risk genes across neuropsychiatric disorders

We derived novel dynamic transcriptomic imputation (dTI) models capable of predicting environmentally dynamic genetically regulated gene expression (GxE-REx) by incorporating genotype-stress interactions across all four postmortem brain regions. These dTI models may be used to predict baseline genetically regulated gene expression (GREx for 11,473-11,965 genes across brain regions, validation R^2^=0.43, 0.44, 0.49, 0.46 for BLA, dACC, DLPFC, MeA, respectively) and stress-interactive genetically regulated gene expression (GxE-Rex for 11,268-11,841 genes across brain regions, validation R^2^=0.45, 0.45, 0.49, 0.46) (**Figure 5A, S. Figure 8A**). Each set of models predicted a subset of genes more accurately (i.e. in the DLPFC, 5,875 genes were predicted more accurately in the stress-interactive model and 6,996 genes were predicted more accurately in the base model) (**Figure 5B**).

**Figure 5:**
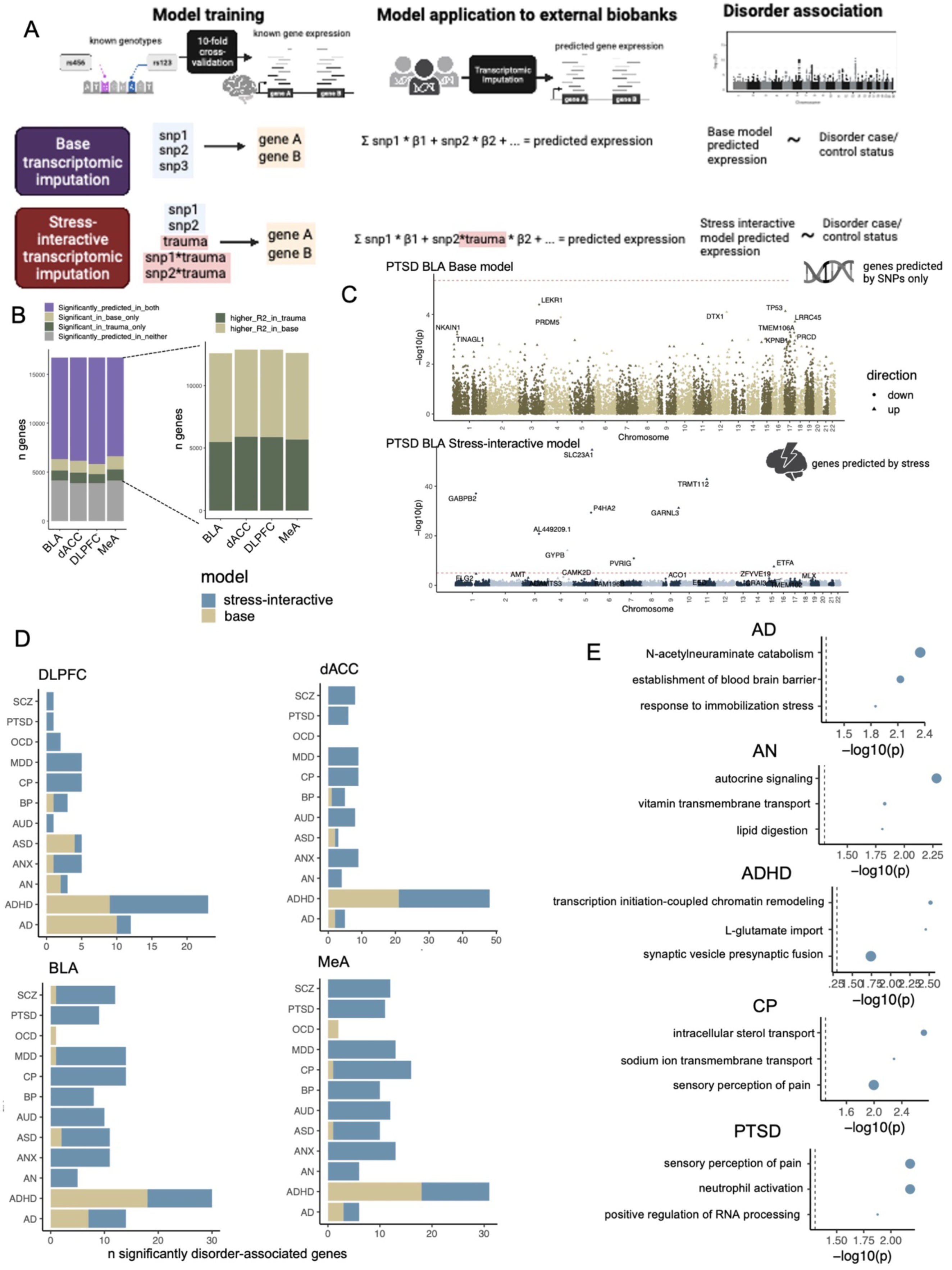
Stress-interactive genetic regulation of expression underlies risk across neuropsychiatric disorders. A) Schematic showing training and validation of base- and stress-interactive transcriptomic imputation models and application to external biobanks to identify genotype-predicted and genotype-and-stress predicted genes associated with neuropsychiatric disorders. B) Bar plot indicating cross-validation predictive accuracy of base and stress-interactive transcriptomic imputation models (left). Of significantly predicted genes in either model, proportion of genes more accurately predicted by base and stress-interactive models (right). C) Manhattan plot showing transcriptome-wide association results of imputed differential expression by the base (top) and stress-interactive (bottom) model as applied to PTSD. Red dotted line indicates genome-wide significance. D) Number of significantly disorder-associated genes identified by transcriptome-wide association studies for 12 neuropsychiatric traits by base (gold) and stress-interactive (blue) models in each post-mortem brain region. Inserts show proportion of genes identified by stress-interactive models with an underlying most significant eQTL in a particular cell type. E) Gene ontology enrichments of stress-interactive model-predicted disorder-associated genes for AD=Alzheimer’s disease, AN=anorexia nervosa, ADHD=attention-deficit hyperactivity disorder, CP=chronic pain, PTSD=post-traumatic stress disorder. Dotted black line indicates significance.

Traumatic stress exacerbates risk for neuropsychiatric disorders beyond PTSD. application of these novel stress-interactive models to 12 neuropsychiatric disorders in the UK Biobank (n=157,322) and Mount Sinai BioMe Biobank (n=28,250) (**Figure 5A**) identified 124 novel psychiatric and 15 novel neurodegenerative GxE-REx associations; the implication is that these genes confer risk only in the context of traumatic stress (**Figure 5D**). For example, nine genes were identified with significant associations (p<1×10^-5^) between predicted BLA GxE-REx and PTSD case/control status only when accounting for traumatic stress (*GABPB2, SLC23A1, AL449209.1, GYPB, P4HA2, PVRIG, GARNL3, TRMT112, ETFA*); no associations were identified using the base models (**Figure 5C**). In Alzheimer’s disease (AD), the BLA base model identified seven AD-associated genes (*DTX1, C3orf37, MEX3C, INPP1, USP32, YPEL2, SENP3)* while the stress-aware model identified six additional genes (*OR52N4, UBXN2B, TDGF1, RBBP6, CBX3, C1orf65*) (**S. Figure 8B**).

Novel brain disorder-associated genes identified by stress-interactive models were enriched in pathways associated previously with stress-dependent disorder risk, such as N-acetylneuraminate catabolism in Alzheimer’s disease (p=4.48×10^-3^), previously implicated in immune-mediated memory deficits^85^, nutrient absorption and metabolism (vitamin transmembrane transport, p=1.49×10^-2^, lipid digestion, p=1.56×10^-2^) in anorexia nervosa, synaptic transmission (L-glutamate import, p=3.5×10^-3^, presynaptic vesicle fusion, p=1.81×10^-2^) in ADHD^86^, and sensory perception of pain^87^ in PTSD^88^ (p=6.66×10^-3^) and chronic pain (p=1.01×10^-2^) (**Figure 5E**).

## DISCUSSION

We demonstrate genotype- and stress-dependent transcriptomic effects detectable long after an initial traumatic exposure. Prompted in part by our observation that hiPSC-derived neurons exhibit intrinsic differential transcriptomic susceptibility to glucocorticoid-induced stress^31^, we examined the genotypic contribution to differential transcriptional encoding of traumatic stress. Common genetic variants that alter transcriptional responses to traumatic stress were identified in post-mortem brains (n=304) and were replicated in an hiPSC-derived neuronal model of glucocorticoid exposure (n=40), indicating together not only that neuronal stress responses are consistent between *in vivo* and *in vitro* paradigms, but also seemingly conserved across neurodevelopment. Without consideration of stress, genotype did not explain variance in expression of stress-interactive eGenes (i.e. rs11586632, p=0.11, **Figure 1D**); it is only when considering both genotype and traumatic burden that the eQTL effect emerges (interaction p=3.11e-6, **Figure 1B**), demonstrating regulation of gene expression in a genotype- and stress-dependent manner. Findings in both cohorts converged on variants lying in transcription factor binding sites, such as GR, NFkB, and YY1. Genotype-dependent transcriptional responses to stress were cell type- and brain region-specific, implicating GABAergic pathways of memory consolidation and novel oligodendrocyte, microglial, and endothelial contributions to blood brain barrier integrity and myelination. Moreover, this genetically regulated response to traumatic stress is relevant across neuropsychiatric disorders, implicating genes involved in immune, metabolic, blood brain barrier and synaptic mechanisms.

GWAS have examined the complex genetic risk architecture underlying PTSD, most recently identifying 81 loci significantly associated with risk across the genome^89^, explaining only 5.32% of heritability^89^, consistent with a modification of genetic risk by environmental exposures (e.g., stress)^90^. Likewise, post-mortem brain studies (e.g. *ELFN1)*^42,73^ and transcription-wide association studies (TWAS) (e.g. *SNRNP35)*^91^ identify genes associated with PTSD at baseline. Here, *in vivo* and *in vitro* approaches indicate that convergent mechanisms underlie long-term encoding of stress exposure. The modification of YY1 transcription factor binding sites by variants conferring genotype-dependent stress response, combined with the blunting of glucocorticoid-mediated transcriptional response via knockdown of YY1, suggest that it is a causal mediator of differential molecular responses to stress. Towards this, YY1 is a crucial factor in the development and function of the central nervous system^92^ known to play a role in stress-sensitivity^93^. Functional disruption of a YY1 binding site by the ADHD-associated SNP rs2271338 mediates genotype-dependent neurodevelopmental impacts via downregulation of the ADGRL3 gene^94^; consistent with YY1 underlying hyper-responsivity of gene targets in PTSD to glucocorticoid-induced stress^31^.

Non-neuronal cells, particularly microglia, endothelial cells, and oligodendrocytes, were enriched for genotype-dependent molecular encoding of stress, consistent with the neuroimmune hypothesis of stress-related brain adaptations. Moreover, blood brain barrier integrity, a known mediator of cognition that when disrupted in associated with cognitive decline^95^, may also contribute to genotype-dependent stress response^96^. Genotype-dependent stress-induced disruption to blood brain barrier permeability may mediate stress-induced cognitive decline across neuropsychiatric disorders.

Given that the major measure of traumatic stress exposure used herein was cumulative trauma burden, summing each traumatic experience across the lifespan, a limitation of our analysis is the equal weighting of diverse stressors known to impart different risks for PTSD (i.e., witnessing an accident confers lower risk for PTSD compared to interpersonal traumas such as sexual violence^97^, but are nonetheless weighted equally in this analysis). Likewise, cumulative trauma burden does not discriminate between temporal exposures to traumatic stress (i.e., childhood vs adulthood exposures^98^, or chronic vs acute stress^99^) that are known to differentially contribute to PTSD susceptibility, suggesting differential molecular encoding during critical periods. In the future, weighting and partitioning traumas based on type and severity, coupled with differentiating between childhood and adulthood exposures, may elucidate unique encoding mechanisms for specific types of traumas. Our analyses assumed that multiple traumas linearly impact molecular response, but trauma burden may in fact confer a nonlinear effect^7,100^. Studies examining the subjective experience of trauma are necessary to discern these biological consequences.

hiPSC-derived models present substantial advantages in that they model the impact of a controlled biological stressor, therefore permitting isogenic comparisons of pre- and post-stress states of transcriptomic regulation. This allows for assessment of causal glucocorticoid-induced regulatory changes, rather than associations confounded by diverse lifetime experiences and varied donor genetic backgrounds. Nonetheless, they are limited in that they model only a single aspect of stress, without consideration of other physiological mediators (e.g., catecholamine reactivity^101^, sympathetic cholinergic activation^102^, and pro-inflammatory cytokines^103^). Moreover, the glucocorticoid-stimulated neurons studied herein approximate acute encoding of stress, whereas post-mortem brain signatures likely also embody aspects of recovery and response to stress. *In vitro* experiments assessing acute stress withdrawal may distinguish encoding of stress exposure from encoding of stress recovery.

Assessed in their natural contexts, the effects of genetic variants may have been confounded by other variants in high linkage disequilibrium, potentially obscuring true causal variants. Massively parallel reporter assays^104,105^ (MPRAs) applied to assess allele-dependent transcriptional activity under glucocorticoid exposure contexts^106^ could empirically resolve true causal variants underlying genotype-dependent stress encoding. Likewise, expansion of CRISPR screens of non-coding variants (e.g., CRISPR-QTL^107^) and/or analysis of pools of dozens of donors (e.g., village-in-a-dish^108^) may pinpoint putative enhancers with stress-dependent regulatory activity and reveal their downstream target genes.

While traditional functional annotation of brain (or neuropsychiatric) disorder-associated variants does not account for environmental impacts^109^, here we demonstrate that eQTLs are sensitive to environmental interactions. Thus, genetics-only approaches to brain disorder biology fail to capture regulatory mechanisms associated with risk under certain environmental contexts, missing important disorder-associated variants, genes, and pathways. Integrating environmental interactions across a variety of brain disorder-associated contexts to eQTL studies and their application to functionally annotate GWAS variants will likely uncover novel pathways of gene x environment interactions essential to brain disorder biology, informing mechanisms by which the impact of genetic risk can be modified. These pathways are likely to have clear translational value in diagnosing, preventing, or treating disease. First, future polygenic risk scores should include environmental measures of traumatic stress to stratify high-risk individuals more accurately. Second, interventions aimed at reducing neuropsychiatric risk should consider mitigating the exposure to and impact of traumatic experiences on particularly vulnerable individuals. Given that diverse environmental exposures may have distinct molecular encoding effects, we urge expansion of this work to broadly consider all gene x environment interactions linked to brain disorder diagnosis and outcomes (e.g., infection, toxins, drugs of addiction, medications), to better understand disorder incidence and treatment response.

Biological vulnerability to stress is dependent on both inherent genotype and the extent of traumatic exposure. Variants conferring differential susceptibility to traumatic stress likely broadly confer neuropsychiatric disorder risk far beyond PTSD, necessitating consideration of the impact of lifetime trauma across brain traits, disorders, and diseases. The cross-disorder relevance of stress-interactive variants underscores the importance of collecting detailed trauma histories clinically, even in patients not deemed otherwise biologically vulnerable, as traumatic stress may confer risk through novel mechanisms. Finally, if stress and trauma indeed result in long-term encoding of cross-disorder risk, this suggests a convergent point of therapeutic intervention to increase resilience, improve brain health, and prevent disease.

## Supplemental Author List

NYSCF Global Array Team: Lauren Bauer, Katie Brenner, Geoff Buckley-Herd, Sean DesMarteau, Patrick Fenton, Peter Ferrarotto, Jenna Hall, Selwyn Jacob, Travis Kroeker, Gregory Lallos, Hector Martinez, Paul McCoy, Frederick J. Monsma, Dorota Moroziewicz, Reid Otto, Kathryn Reggio, Bruce Sun, Rebecca Tibbets, Dong Woo Shin, Hongyan Zhou & Matthew Zimmer Traumatic Stress Brain Research Group: Victor E. Alvarez, David Benedek, Alicia Che, Dianne A. Cruz, David A. Davis, Matthew J. Girgenti, Ellen Hoffman, Paul E. Holtzheimer, Bertrand R. Huber, Alfred Kaye, John H. Krystal, Adam T. Labadorf, Terence M. Keane, Mark W. Logue, Ann McKee, Brian Marx, Mark W. Miller, Crystal Noller, Janitza Montalvo-Ortiz, Meghan Pierce, William K. Scott, Paula Schnurr, Krista DiSano, Thor Stein, Robert Ursano, Douglas E. Williamson, Erika J. Wolf, Keith A. Young, PhD

## AUTHOR CONTRIBUTIONS

Study conception was provided by L.M.H, K.J.B, and C.S. Neuronal generation and modeling was provided by P.J.M.D, T.R., B.M, S.D, J.G, S.A.N, D.P. Data analysis and/or code was provided by C.S, R.S, E.M.H, H.Y, A.C, K.T, C.X. All authors interrelated data and critically revised the manuscript for important intellectual content. hiPSC cohort data and specimen collections were carried out by H.B. and R.Y. Post-mortem cohort data and specimen collections were carried out by M.J.G, J.H.K, P.E.H, K.A.Y, D.B. All authors wrote and approved the final manuscript.

## ACKNOWLEDGEMENTS

This work was supported in part through the computational and data resources and staff expertise provided by Scientific Computing and Data at the Icahn School of Medicine at Mount Sinai and supported by the Clinical and Translational Science Awards (CTSA) grant UL1TR004419 from the National Center for Advancing Translational Sciences. Research reported in this publication was also supported by the Office of Research Infrastructure of the National Institutes of Health under award number S10OD026880 and S10OD030463. The content is solely the responsibility of the authors and does not necessarily represent the official views of the National Institutes of Health. This work was supported with resources and use of facilities at the VA Connecticut Health Care System, West Haven, CT. This work was funded in part by the State of Connecticut, Department of Mental Health and Addiction Services. The views expressed here are those of the authors and do not necessarily reflect the position or policy of the US Department of Veterans Affairs (VA) or the U.S. government or the views of the Department of Mental Health and Addiction Services or the State of Connecticut.

## Supplementary Figures

**Figure S1.**
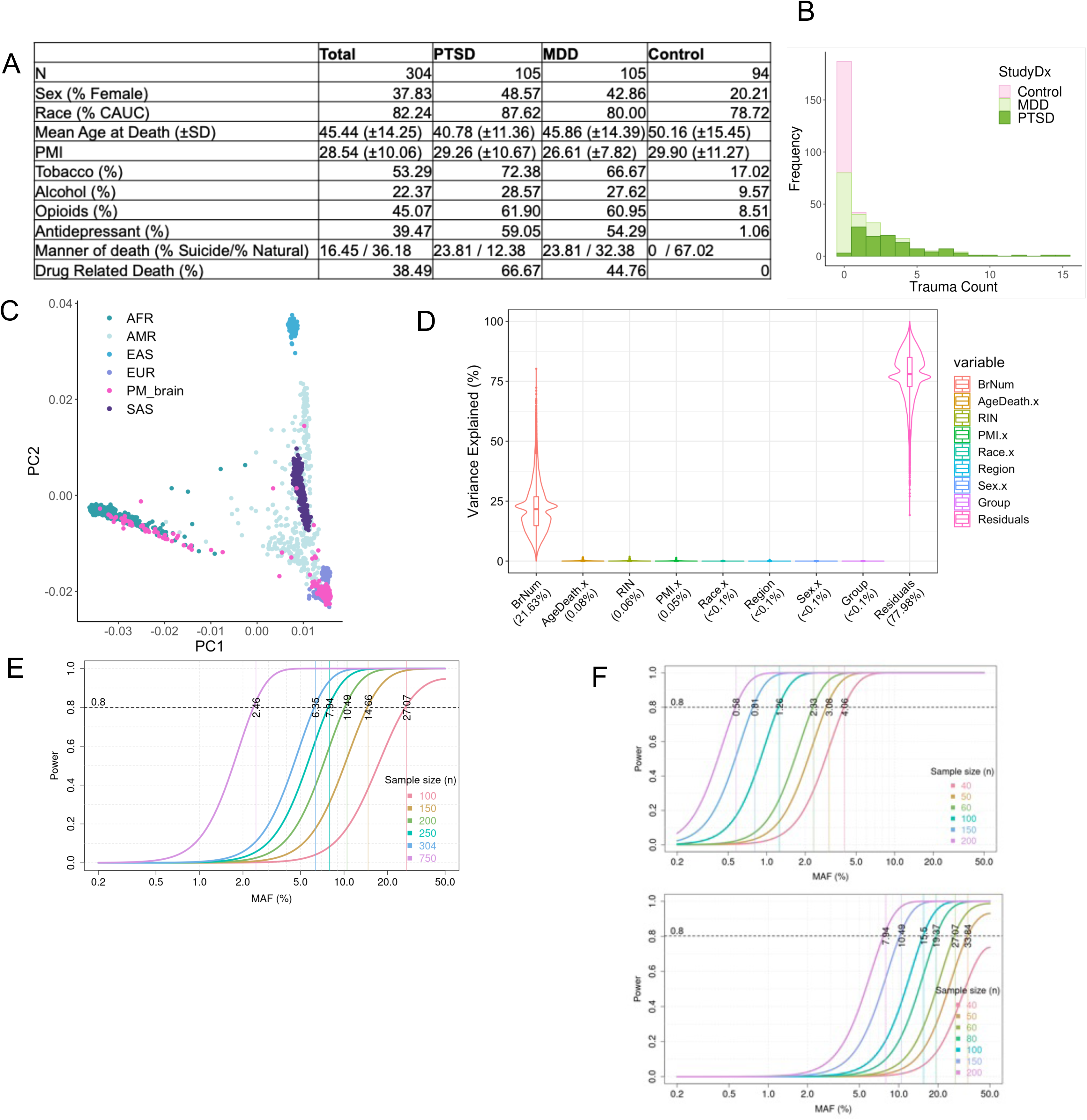
Cohort and power descriptions. A) Post-mortem brain cohort demographics split by diagnosis. B) Frequency of cumulative traumatic exposures in post-mortem brain diagnosis, colored by diagnosis. C) Genotype-derived principal components projected onto the 1000 genomes cohort indicative of ancestry mapping. D) VariancePartition plots indicating percent of variance explained of various covariates in the post-mortem brain cohort. E) Power to detect eQTL effects in the post-mortem brain cohort based on GTEx post-mortem eQTLs. F) Power to detect eQTL effects in the hiPSC-derived neuronal cohort based on assumptions for post-mortem analysis (above) and hiPSC-corrected SD assumption (below)

**Figure S2.**
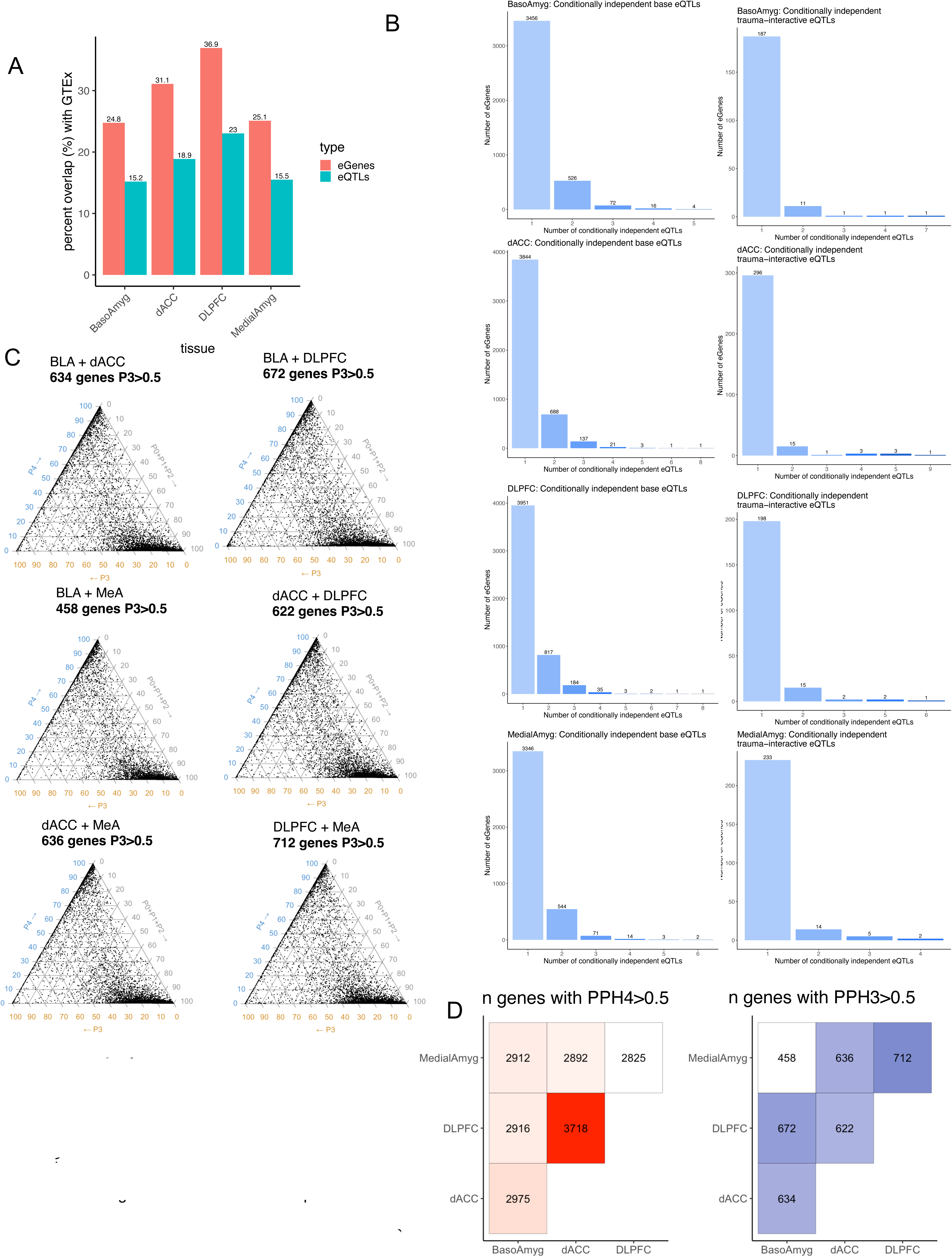
Post mortem brain eQTL replication and region comparison. A) Percent overlap of eQTLs (blue) and eGenes (red) in this study with GTEx eQTLs in the dACC, DLPFC, and amygdala. B) Number of conditionally independent eQTL signals for each base (left) or stress-interactive (right) eGene in each brain region. C) Pair-wise colocalization between brain regions indicating shared eGenes with independent underlying regulation. D) Heatmap showing number of eGenes with colocalized (left) and independent regulation (right) across brain regions.

**Figure S3.**
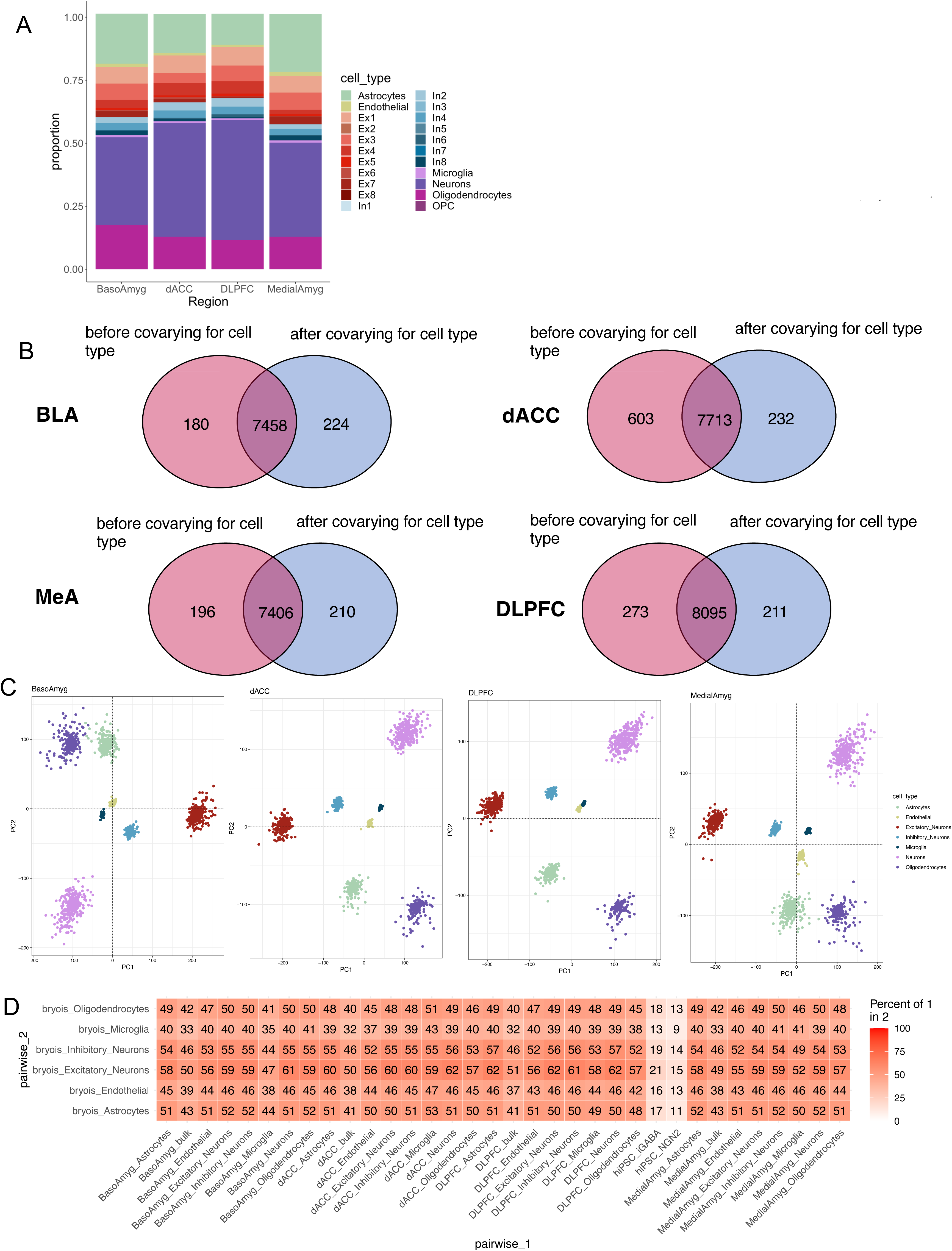
Cell type composition of bulk brain regions. A) Imputed cell type proportions of post-mortem brain regions using psychENCODE cell type reference panels. B) Venn diagram of number of significant eGenes before and after covarying for cell type proportion. C) PCA plots of imputed cell type-specific expression for each brain region. D) Replication of imputed cell type specific eQTLs with previously published single cell eQTLs (Bryois et al)

**Figure S4.**
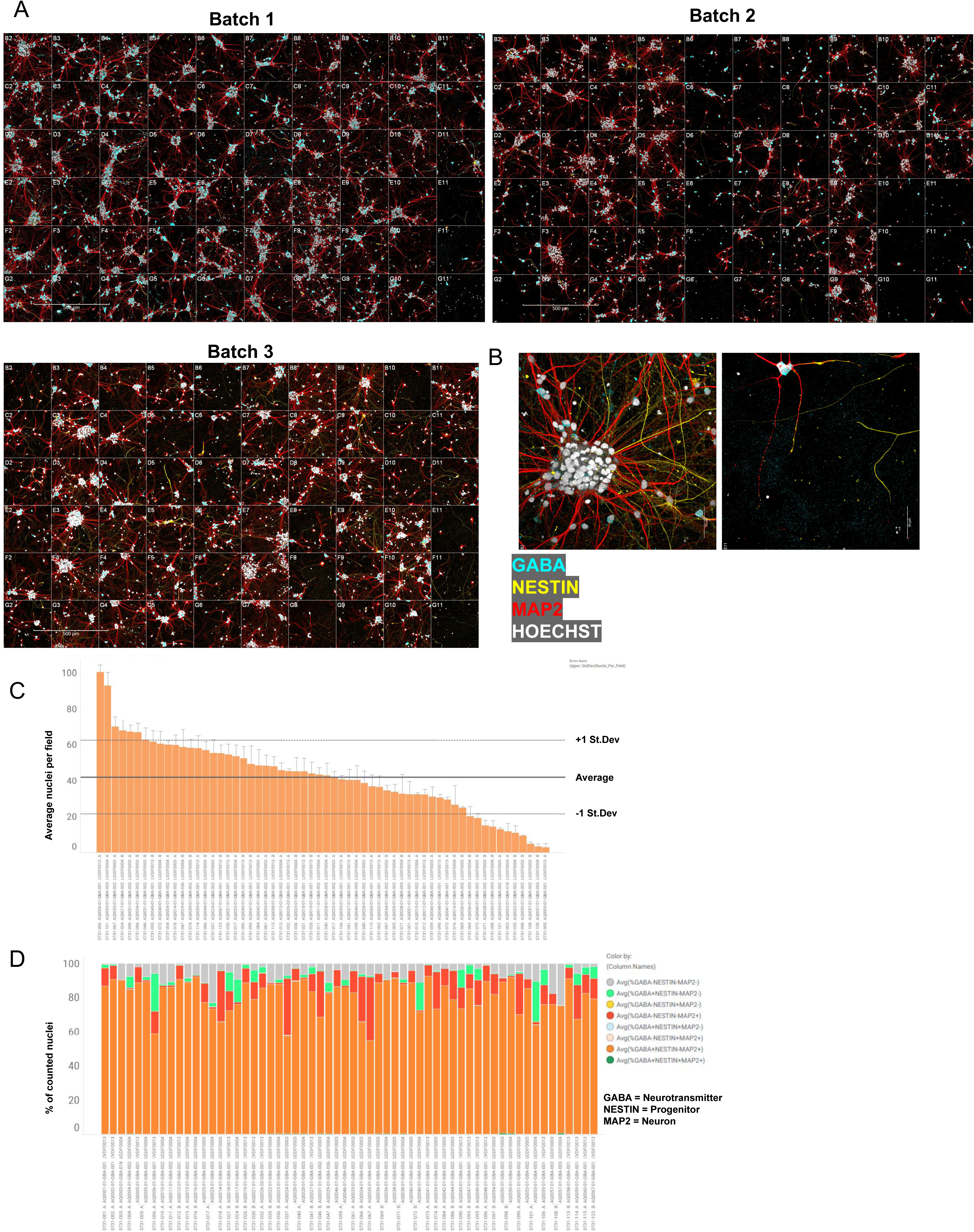
hiPSC-derived GABAergic neurons. (A) Labeled montage of a single 40x field per well. (B) High (AQ0045) and low (AQ0005) density examples enlarged. C) Cell count per field per individual, D) GABA, Progenitor, and Neuron percent per well per individual

**Figure S5.**
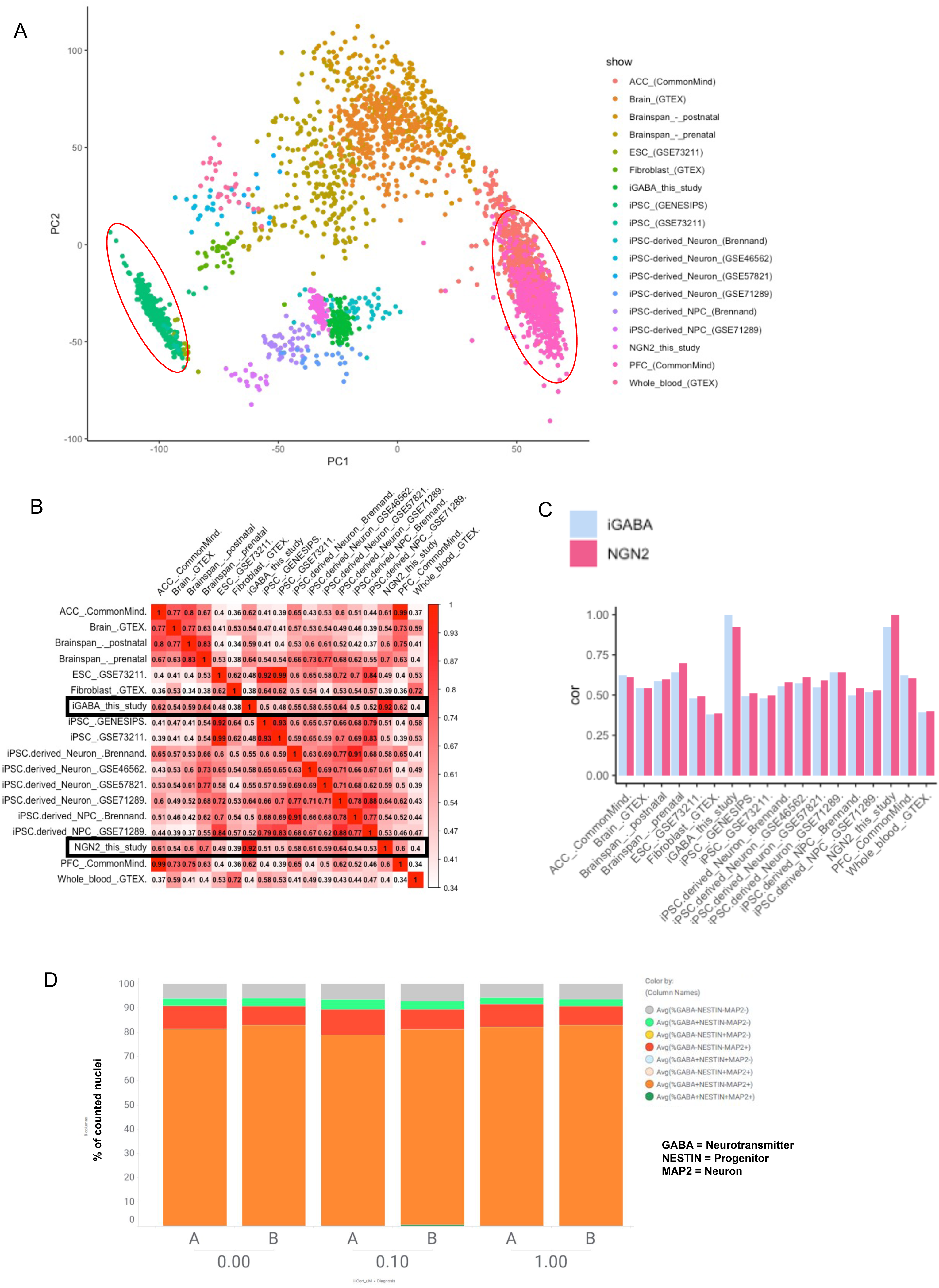
Developmental specificity of hiPSC-derived GABAergic neurons. (A) PCA of GABAergic expression data compared to previous hiPSC-derived neuronal studies, including the excitatory neuron study used here. B) Pairwise correlation between expression signatures of iGABA and iGLUT neurons from our study with cell types across 16 independent studies. C) Correlation between expression signatures of iGABA and iGLUT neurons from our study with other hiPSC studies. D) Proportion of GABA+, NESTIN+, and MAP2+ cells are stable across HCort dosages.

**Figure S6.**
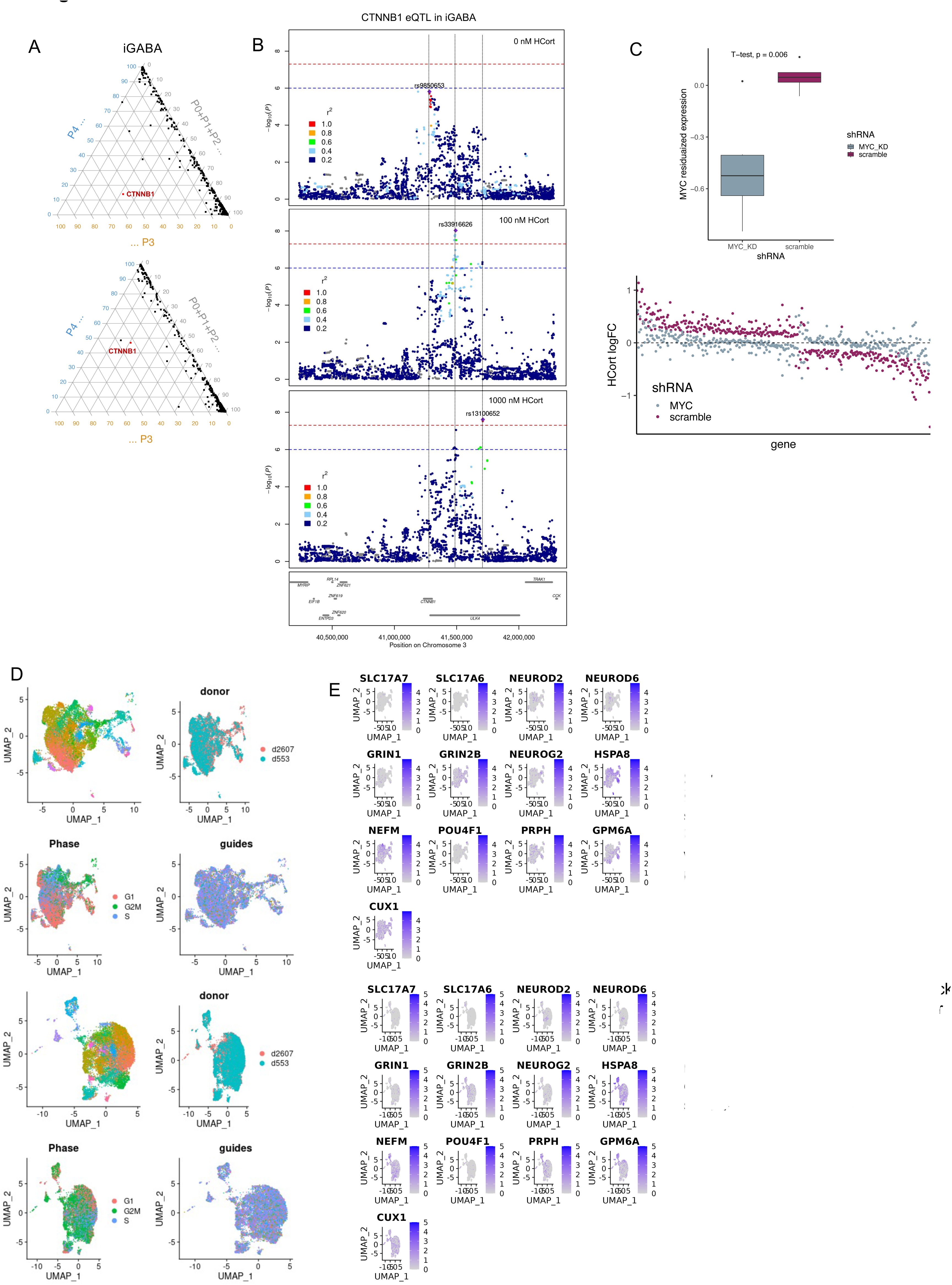
hiPSC validation of stress-interactive eQTLs. (A) Ternary plot showing pair-wise colocalization of HCort dosages. CTNNB1 highlighted with independent underlying regulation (PPH3>0.5). B) CTNNB1 LocusZoom showing eQTL signal at 0nM HCort (top) 100 nM HCort (middle), and 1000 nM HCort (bottom), with three independent peaks regulating expression at each dose. C) (top) shRNA mediated knock down of the MYC transcription factor compared to scramble. (Bottom) The log2FC of HCort-stimulation in HCort-responsive genes in the scramble condition (purple) and with MYC shRNA-mediated knockdown (blue).

**Figure S7.**
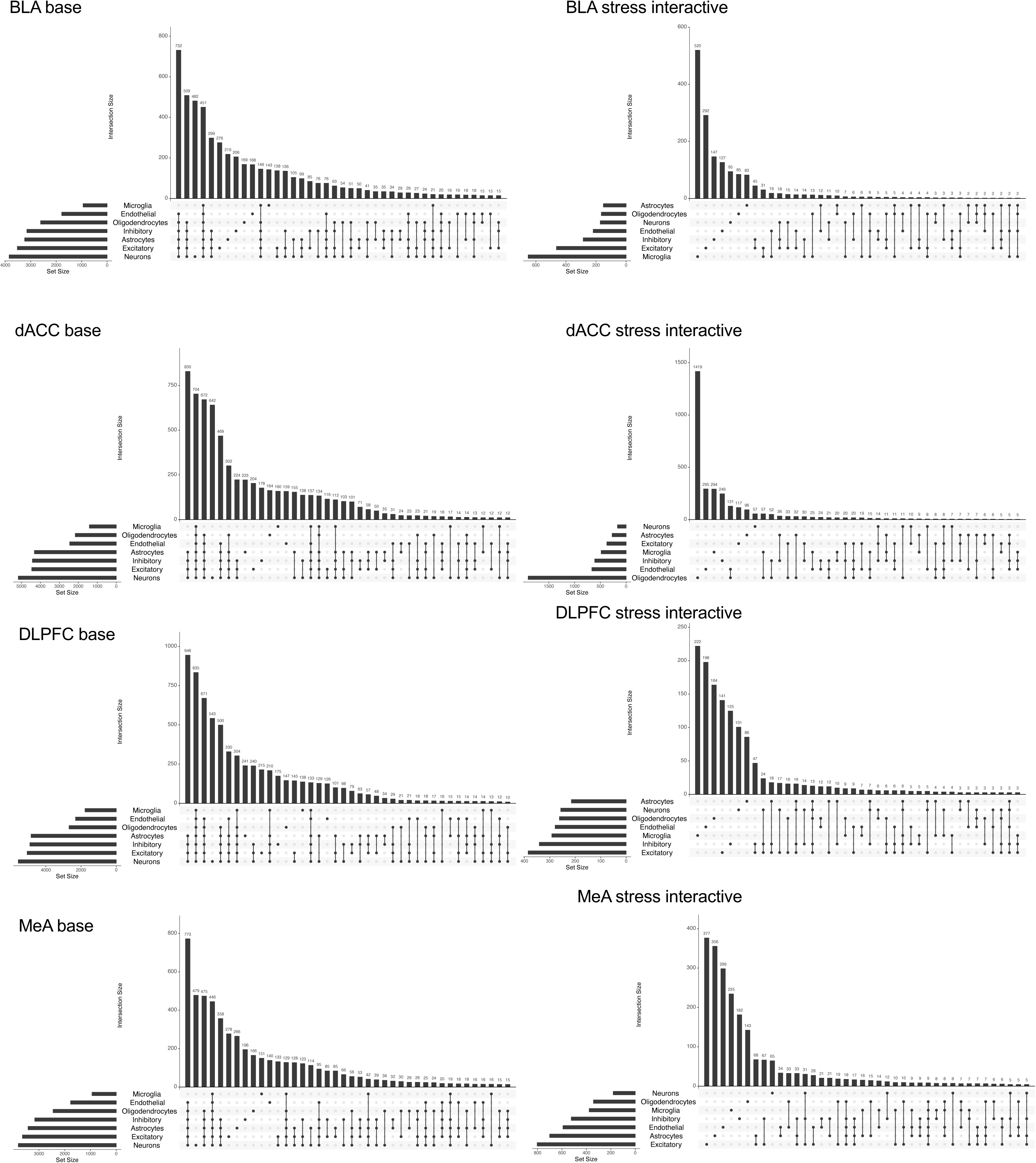
Overlap of cell type imputed eQTLs across the post-mortem brain. UpSet plots indicating overlap of eGenes detected across imputed cell types for base eGenes and stress-interactive eGenes across post mortem brain regions.

**Figure S8.**
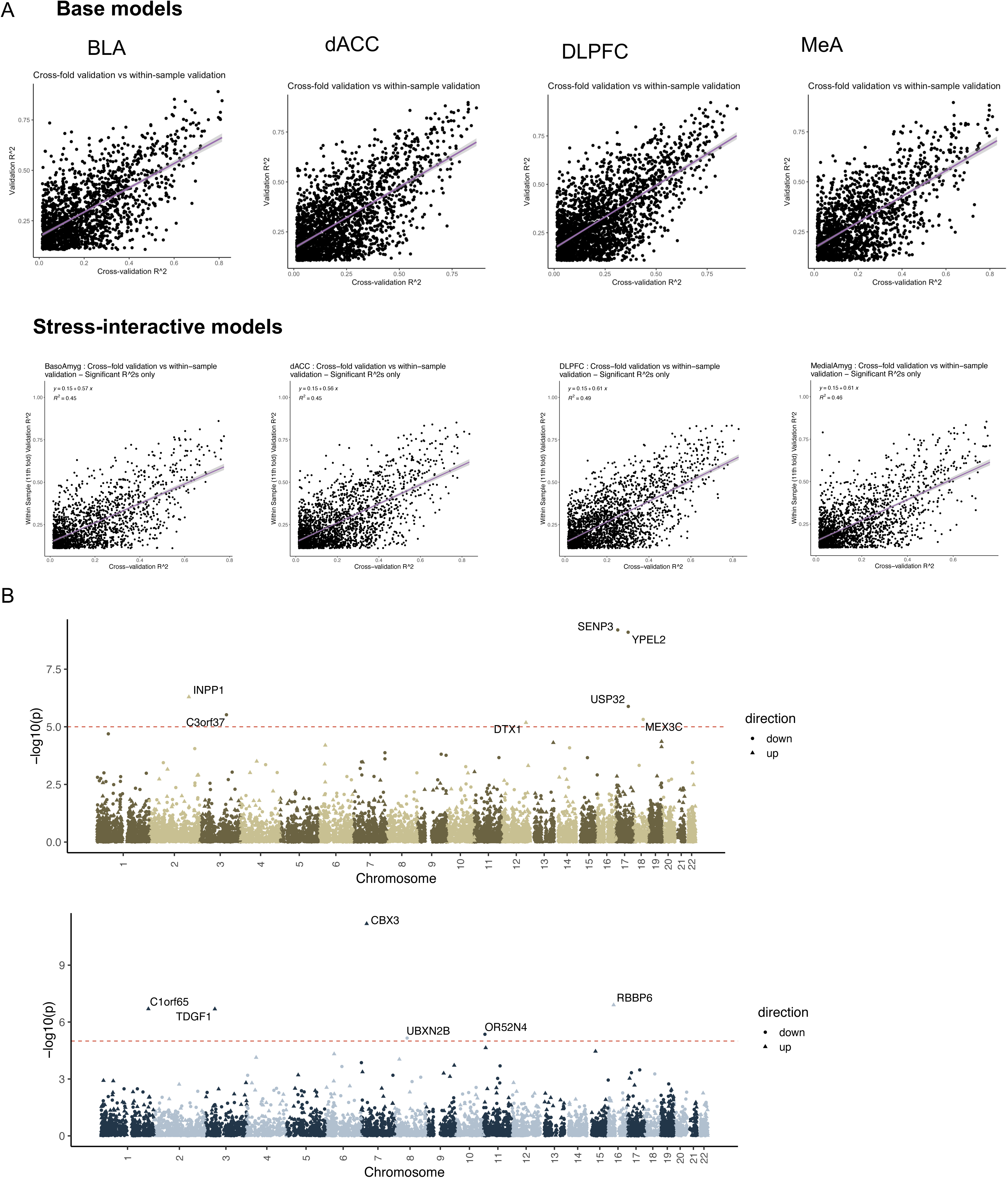
Application of transcriptomic imputation models. A) Cross fold validation R^2^ compared to hold-out fold validation R^2^ for each significantly predicted gene by base transcriptomic imputation (above) and stress-interactive transcriptomic imputation models (below). B) Manhattan plot showing transcriptome-wide association results of imputed differential expression by the base (top) and stress-interactive (bottom) model as applied to Alzheimer’s disease. Red dotted line indicates genome-wide significance.

## REFERENCES

1. McFARLANE, A. C. The long-term costs of traumatic stress: intertwined physical and psychological consequences. World Psychiatry 9, 3–10 (2010).

2. Breslau, N., Davis, G. C., Peterson, E. L. & Schultz, L. R. A second look at comorbidity in victims of trauma: the posttraumatic stress disorder–major depression connection. Biol. Psychiatry 48, 902–909 (2000).

3. Breslau, N., Davis, G. C. & Schultz, L. R. Posttraumatic Stress Disorder and the Incidence of Nicotine, Alcohol, and Other Drug Disorders in Persons Who Have Experienced Trauma. Arch. Gen. Psychiatry 60, 289–294 (2003).

4. McLaughlin, K. A. et al. Childhood Adversities and First Onset of Psychiatric Disorders in a National Sample of US Adolescents. Arch. Gen. Psychiatry 69, 1151–1160 (2012).

5. LeMoult, J. et al. Meta-analysis: Exposure to Early Life Stress and Risk for Depression in Childhood and Adolescence. J. Am. Acad. Child Adolesc. Psychiatry 59, 842–855 (2020).

6. Aas, M. et al. The role of childhood trauma in bipolar disorders. Int. J. Bipolar Disord. 4, 2 (2016).

7. Marchese, S. & Huckins, L. M. Trauma Matters: Integrating Genetic and Environmental Components of PTSD. Adv. Genet. Hoboken NJ 4, 2200017 (2023).

8. Justice, N. J. The relationship between stress and Alzheimer’s disease. Neurobiol. Stress 8, 127–133 (2018).

9. Popovic, D. et al. Childhood Trauma in Schizophrenia: Current Findings and Research Perspectives. Front. Neurosci. 13, 274 (2019).

10. Tagay, S., Schlottbohm, E., Reyes-Rodriguez, M. L., Repic, N. & Senf, W. Eating Disorders, Trauma, PTSD and Psychosocial Resources. Eat. Disord. 22, 33–49 (2014).

11. Hoppen, T. H. & Morina, N. The prevalence of PTSD and major depression in the global population of adult war survivors: a meta-analytically informed estimate in absolute numbers. Eur. J. Psychotraumatology 10, 1578637 (2019).

12. Feder, A., Nestler, E. J. & Charney, D. S. Psychobiology and molecular genetics of resilience. Nat. Rev. Neurosci. 10, 446–457 (2009).

13. Bremner, J. D. Traumatic stress: effects on the brain. Dialogues Clin. Neurosci. 8, 445 (2006).

14. Finsterwald, C. & Alberini, C. M. Stress and glucocorticoid receptor-dependent mechanisms in long-term memory: from adaptive responses to psychopathologies. Neurobiol. Learn. Mem. 0, 17–29 (2014).

15. Laine, M. A. & Shansky, R. M. Rodent models of stress and dendritic plasticity – Implications for psychopathology. Neurobiol. Stress 17, 100438 (2022).

16. Russo, S. J., Murrough, J. W., Han, M.-H., Charney, D. S. & Nestler, E. J. Neurobiology of resilience. Nat. Neurosci. 15, 1475–1484 (2012).

17. Goldwater, D. S. et al. Structural and functional alterations to rat medial prefrontal cortex following chronic restraint stress and recovery. Neuroscience 164, 798–808 (2009).

18. Chang, C. & Grace, A. A. Amygdala-Ventral Pallidum Pathway Decreases Dopamine Activity After Chronic Mild Stress in Rats. Biol. Psychiatry 76, 223–230 (2014).

19. Liston, C. et al. Stress-induced alterations in prefrontal cortical dendritic morphology predict selective impairments in perceptual attentional set-shifting. J. Neurosci. 26, 7870–7874 (2006).

20. Liston, C. & Gan, W. B. Glucocorticoids are critical regulators of dendritic spine development and plasticity in vivo. Proc Natl Acad Sci USA 108, (2011).

21. Peña, C. J. et al. Early life stress alters transcriptomic patterning across reward circuitry in male and female mice. Nat. Commun. 10, 5098 (2019).

22. Smoller, J. W. The Genetics of Stress-Related Disorders: PTSD, Depression, and Anxiety Disorders. Neuropsychopharmacology 41, 297–319 (2016).

23. Caspi, A. et al. Influence of life stress on depression: moderation by a polymorphism in the 5-HTT gene. Science 301, 386–389 (2003).

24. Caspi, A. et al. Role of genotype in the cycle of violence in maltreated children. Science 297, 851–854 (2002).

25. Binder, E. B. et al. Polymorphisms in FKBP5 are associated with increased recurrence of depressive episodes and rapid response to antidepressant treatment. Nat. Genet. 36, 1319–1325 (2004).

26. Binder, E. B. et al. Association of FKBP5 polymorphisms and childhood abuse with risk of posttraumatic stress disorder symptoms in adults. JAMA 299, 1291–1305 (2008).

27. Fani, N. et al. FKBP5 and attention bias for threat: associations with hippocampal function and shape. JAMA Psychiatry 70, 392–400 (2013).

28. Klengel, T. et al. Allele-specific FKBP5 DNA demethylation mediates gene–childhood trauma interactions. Nat. Neurosci. 16, 33–41 (2013).

29. Elbau, I. G., Cruceanu, C. & Binder, E. B. Genetics of Resilience: Gene-by-Environment Interaction Studies as a Tool to Dissect Mechanisms of Resilience. Biol. Psychiatry 86, 433– 442 (2019).

30. DePierro, J., Lepow, L., Feder, A. & Yehuda, R. Translating Molecular and Neuroendocrine Findings in Posttraumatic Stress Disorder and Resilience to Novel Therapies. Biol. Psychiatry 86, 454–463 (2019).

31. Seah, C. et al. Modeling gene × environment interactions in PTSD using human neurons reveals diagnosis-specific glucocorticoid-induced gene expression. Nat. Neurosci. 25, 1434–1445 (2022).

32. Fu, J. et al. Unraveling the Regulatory Mechanisms Underlying Tissue-Dependent Genetic Variation of Gene Expression. PLOS Genet. 8, e1002431 (2012).

33. Aguet, F. et al. Genetic effects on gene expression across human tissues. Nature 550, 204– 213 (2017).

34. Moore, S. R., et al. Sex differences in the genetic regulation of the blood transcriptome response to glucocorticoid receptor activation. 2020.10.19.20213983 https://www.medrxiv.org/content/10.1101/2020.10.19.20213983v3 (2021) doi:10.1101/2020.10.19.20213983.

35. Yao, C. et al. Sex- and age-interacting eQTLs in human complex diseases. Hum. Mol. Genet. 23, 1947–1956 (2014).

36. Lindén, M. et al. Sex influences eQTL effects of SLE and Sjögren’s syndrome-associated genetic polymorphisms. Biol. Sex Differ. 8, 34 (2017).

37. Werling, D. M. et al. Whole-Genome and RNA Sequencing Reveal Variation and Transcriptomic Coordination in the Developing Human Prefrontal Cortex. Cell Rep. 31, 107489 (2020).

38. Cuomo, A. S. E. et al. Single-cell RNA-sequencing of differentiating iPS cells reveals dynamic genetic effects on gene expression. Nat. Commun. 11, 810 (2020).

39. Knowles, D. A. et al. Determining the genetic basis of anthracycline-cardiotoxicity by molecular response QTL mapping in induced cardiomyocytes. eLife 7, e33480 (2018).

40. Nakagawa, S. et al. Effects of post-traumatic growth on the dorsolateral prefrontal cortex after a disaster. Sci. Rep. 6, 34364 (2016).

41. Alexandra Kredlow, M., Fenster, R. J., Laurent, E. S., Ressler, K. J. & Phelps, E. A. Prefrontal cortex, amygdala, and threat processing: implications for PTSD. Neuropsychopharmacology 47, 247–259 (2022).

42. Jaffe, A. E. et al. Decoding Shared Versus Divergent Transcriptomic Signatures Across Cortico-Amygdala Circuitry in PTSD and Depressive Disorders. Am. J. Psychiatry 179, 673– 686 (2022).

43. Liberzon, I. & Sripada, C. S. The functional neuroanatomy of PTSD: a critical review. in Progress in Brain Research (eds. De Kloet, E. R., Oitzl, M. S. & Vermetten, E.) vol. 167 151–169 (Elsevier, 2007).

44. Koek, R. J. et al. Deep brain stimulation of the basolateral amygdala for treatment-refractory combat post-traumatic stress disorder (PTSD): study protocol for a pilot randomized controlled trial with blinded, staggered onset of stimulation. Trials 15, 356 (2014).

45. Zhang, H.-H. et al. Traumatic Stress Produces Delayed Alterations of Synaptic Plasticity in Basolateral Amygdala. Front. Psychol. 10, 2394 (2019).

46. Huckins, L. M., et al. Polygenic regulation of PTSD severity and outcomes among World Trade Center responders. 2020.12.06.20244772 https://www.medrxiv.org/content/10.1101/2020.12.06.20244772v2 (2021) doi:10.1101/2020.12.06.20244772.

47. Huckins, L., Pietrzak, R., Yehuda, R., Stahl, E. & Feder, A. Genetic Regulation of PTSD and Resilience Among World Trade Center Responders. Biol. Psychiatry 87, S26 (2020).

48. Marchese, S. et al. Altered gene expression and PTSD symptom dimensions in World Trade Center responders. Mol. Psychiatry 27, 2225–2246 (2022).

49. Barretto, N. et al. ASCL1- and DLX2-induced GABAergic neurons from hiPSC-derived NPCs. J. Neurosci. Methods 334, 108548 (2020).

50. Lonsdale, J. et al. The Genotype-Tissue Expression (GTEx) project. Nat. Genet. 45, 580– 585 (2013).

51. Schrode, N. et al. Synergistic effects of common schizophrenia risk variants. Nat. Genet. 51, 1475–1485 (2019).

52. DeBoever, C. et al. Large-Scale Profiling Reveals the Influence of Genetic Variation on Gene Expression in Human Induced Pluripotent Stem Cells. Cell Stem Cell 20, 533–546.e7 (2017).

53. Hoffman, G. E. et al. CommonMind Consortium provides transcriptomic and epigenomic data for Schizophrenia and Bipolar Disorder. Sci. Data 6, 180 (2019).

54. Dobbyn, A. et al. Landscape of Conditional eQTL in Dorsolateral Prefrontal Cortex and Co-localization with Schizophrenia GWAS. Am. J. Hum. Genet. 102, 1169–1184 (2018).

55. Giambartolomei, C. et al. Bayesian test for colocalisation between pairs of genetic association studies using summary statistics. PLoS Genet. 10, e1004383 (2014).

56. Watanabe, K., Taskesen, E., van Bochoven, A. & Posthuma, D. Functional mapping and annotation of genetic associations with FUMA. Nat. Commun. 8, 1826 (2017).

57. Newman, A. M. et al. Determining cell type abundance and expression from bulk tissues with digital cytometry. Nat. Biotechnol. 37, 773–782 (2019).

58. Darmanis, S. et al. A survey of human brain transcriptome diversity at the single cell level. Proc. Natl. Acad. Sci. U. S. A. 112, 7285–7290 (2015).

59. Lake, B. B. et al. Neuronal subtypes and diversity revealed by single-nucleus RNA sequencing of the human brain. Science 352, 1586–1590 (2016).

60. Lake, B. B. et al. Integrative single-cell analysis of transcriptional and epigenetic states in the human adult brain. Nat. Biotechnol. 36, 70–80 (2018).

61. Bayesian estimation of cell type-specific gene expression with prior derived from single-cell data - PubMed. https://pubmed.ncbi.nlm.nih.gov/33837133/.

62. Leek, J. T., Johnson, W. E., Parker, H. S., Jaffe, A. E. & Storey, J. D. The sva package for removing batch effects and other unwanted variation in high-throughput experiments. Bioinforma. Oxf. Engl. 28, 882–883 (2012).

63. Ward, L. D. & Kellis, M. HaploReg v4: systematic mining of putative causal variants, cell types, regulators and target genes for human complex traits and disease. Nucleic Acids Res. 44, D877–D881 (2016).

64. Bray, N. L., Pimentel, H., Melsted, P. & Pachter, L. Near-optimal probabilistic RNA-seq quantification. Nat. Biotechnol. 34, 525–527 (2016).

65. Korsunsky, I. et al. Fast, sensitive and accurate integration of single-cell data with Harmony. Nat. Methods 16, 1289–1296 (2019).

66. Friedman, J. H., Hastie, T. & Tibshirani, R. Regularization Paths for Generalized Linear Models via Coordinate Descent. J. Stat. Softw. 33, 1–22 (2010).

67. Marchese, S., Cuddleston, W., Seah, C., Johnson, J. & Huckins, L. M. Disentangling the roles of trauma and genetics in psychiatric disorders using an Electronic Health Records-based approach. 2022.08.28.22279306 Preprint at 10.1101/2022.08.28.22279306 (2022).

68. Kheradpour, P. & Kellis, M. Systematic discovery and characterization of regulatory motifs in ENCODE TF binding experiments. Nucleic Acids Res. 42, 2976–2987 (2014).

69. Rao, N. A. S. et al. Coactivation of GR and NFKB alters the repertoire of their binding sites and target genes. Genome Res. 21, 1404–1416 (2011).

70. Gray, J. D., Kogan, J. F., Marrocco, J. & McEwen, B. S. Genomic and epigenomic mechanisms of glucocorticoids in the brain. Nat. Rev. Endocrinol. 13, 661–673 (2017).

71. Huang, J. et al. Involvement of the GABAergic system in PTSD and its therapeutic significance. Front. Mol. Neurosci. 16, 1052288 (2023).

72. Daskalakis, N. P. Cell type-specific Dissection of PTSD and MDD in Human Post-Mortem DLPFC reveals genetic and glucocorticoid regulation. Psychoneuroendocrinology 131, 105547 (2021).

73. Girgenti, M. J. et al. Transcriptomic organization of the human brain in post-traumatic stress disorder. Nat. Neurosci. 24, 24–33 (2021).

74. Ho, S. M. Rapid Ngn2-induction of excitatory neurons from hiPSC-derived neural progenitor cells. Methods 101, (2016).

75. Zhang, Y. Rapid single-step induction of functional neurons from human pluripotent stem cells. Neuron 78, (2013).

76. Flaherty, E. Neuronal impact of patient-specific aberrant NRXN1alpha splicing. Nat Genet 51, (2019).

77. Yang, N. et al. Generation of pure GABAergic neurons by transcription factor programming. Nat. Methods 14, 621–628 (2017).

78. Forrest, M. P. et al. Open Chromatin Profiling in hiPSC-Derived Neurons Prioritizes Functional Noncoding Psychiatric Risk Variants and Highlights Neurodevelopmental Loci. Cell Stem Cell 21, 305–318.e8 (2017).

79. Chavez, A. et al. Highly-efficient Cas9-mediated transcriptional programming. Nat. Methods 12, 326–328 (2015).

80. Gilbertson, M. W. et al. Smaller hippocampal volume predicts pathologic vulnerability to psychological trauma. Nat. Neurosci. 5, 1242–1247 (2002).

81. Bryois, J. et al. Cell-type-specific cis-eQTLs in eight human brain cell types identify novel risk genes for psychiatric and neurological disorders. Nat. Neurosci. 25, 1104–1112 (2022).

82. Nadra, K. et al. Phosphatidic acid mediates demyelination in Lpin1 mutant mice. Genes Dev. 22, 1647–1661 (2008).

83. Blecharz-Lang, K. G. et al. Interleukin 6-Mediated Endothelial Barrier Disturbances Can Be Attenuated by Blockade of the IL6 Receptor Expressed in Brain Microvascular Endothelial Cells. Transl. Stroke Res. 9, 631–642 (2018).

84. Joshi, B. S. & Zuhorn, I. S. Heparan sulfate proteoglycan-mediated dynamin-dependent transport of neural stem cell exosomes in an in vitro blood-brain barrier model. Eur. J. Neurosci. 53, 706–719 (2021).

85. Suzzi, S. et al. N-acetylneuraminic acid links immune exhaustion and accelerated memory deficit in diet-induced obese Alzheimer’s disease mouse model. Nat. Commun. 14, 1293 (2023).

86. Maltezos, S. et al. Glutamate/glutamine and neuronal integrity in adults with ADHD: a proton MRS study. Transl. Psychiatry 4, e373 (2014).

87. Johnston, K. J. A. & Huckins, L. M. Chronic Pain and Psychiatric Conditions. Complex Psychiatry 9, 24–43 (2022).

88. Jenewein, J. et al. Altered Pain Perception and Fear-Learning Deficits in Subjects With Posttraumatic Stress Disorder. J. Pain Off. J. Am. Pain Soc. 17, 1325–1333 (2016).

89. THE BRAINSTORM CONSORTIUM et al. Analysis of shared heritability in common disorders of the brain. Science 360, eaap8757 (2018).

90. Hunter, D. J. Gene-environment interactions in human diseases. Nat. Rev. Genet. 6, 287– 298 (2005).

91. Huckins, L. M. et al. Analysis of Genetically Regulated Gene Expression Identifies a Prefrontal PTSD Gene, SNRNP35, Specific to Military Cohorts. Cell Rep. 31, 107716 (2020).

92. Zurkirchen, L. et al. Yin Yang 1 sustains biosynthetic demands during brain development in a stage-specific manner. Nat. Commun. 10, 2192 (2019).

93. Kwon, D. Y. et al. Neuronal Yin Yang1 in the prefrontal cortex regulates transcriptional and behavioral responses to chronic stress in mice. Nat. Commun. 13, 55 (2022).

94. Martinez, A. F. et al. An Ultraconserved Brain-Specific Enhancer Within ADGRL3 (LPHN3) Underpins Attention-Deficit/Hyperactivity Disorder Susceptibility. Biol. Psychiatry 80, 943– 954 (2016).

95. Nation, D. A. et al. Blood-brain barrier breakdown is an early biomarker of human cognitive dysfunction. Nat. Med. 25, 270–276 (2019).

96. Iadecola, C. The Neurovascular Unit Coming of Age: A Journey through Neurovascular Coupling in Health and Disease. Neuron 96, 17–42 (2017).

97. Yehuda, R. Post-traumatic stress disorder. Nat Rev Primer 1, (2015).

98. Cloitre, M. et al. A developmental approach to complex PTSD: Childhood and adult cumulative trauma as predictors of symptom complexity. J. Trauma. Stress 22, 399–408 (2009).

99. Pratchett, L. C. & Yehuda, R. Foundations of posttraumatic stress disorder: Does early life trauma lead to adult posttraumatic stress disorder? Dev. Psychopathol. 23, 477–491 (2011).

100. Yehuda, R. & LeDoux, J. Response Variation following Trauma: A Translational Neuroscience Approach to Understanding PTSD. Neuron 56, 19–32 (2007).

101. Sah, R. & Geracioti, T. Neuropeptide Y and posttraumatic stress disorder. Mol. Psychiatry 18, 646–655 (2013).

102. Belkin, M. R. & Schwartz, T. L. Alpha-2 receptor agonists for the treatment of posttraumatic stress disorder. Drugs Context 4, 212286 (2015).

103. Passos, I. C. et al. Inflammatory markers in post-traumatic stress disorder: a systematic review, meta-analysis, and meta-regression. Lancet Psychiatry 2, 1002–1012 (2015).

104. Townsley, K. G., Brennand, K. J. & Huckins, L. M. Massively parallel techniques for cataloguing the regulome of the human brain. Nat. Neurosci. 1–13 (2020) doi:10.1038/s41593-020-00740-1.

105. McAfee, J. C. et al. Systematic investigation of allelic regulatory activity of schizophrenia-associated common variants. Cell Genomics 3, 100404 (2023).

106. Seah, C., Huckins, L. M. & Brennand, K. J. Stem Cell Models for Context-Specific Modeling in Psychiatric Disorders. Biol. Psychiatry 93, 642–650 (2023).

107. Gasperini, M. et al. A Genome-wide Framework for Mapping Gene Regulation via Cellular Genetic Screens. Cell 176, 377–390.e19 (2019).

108. Neavin, D. R. et al. A village in a dish model system for population-scale hiPSC studies. Nat. Commun. 14, 3240 (2023).

109. Young, H., Cote, A. & Huckins, L. M. Chapter 14 - Integration with systems biology approaches and -omics data to characterize risk variation. in Psychiatric Genomics (eds. Tsermpini, E. E., Alda, M. & Patrinos, G. P.) 289–315 (Academic Press, (2022). doi:10.1016/B978-0-12-819602-1.00017-6.

